# Unexpected Complexity of the Ammonia Monooxygenase in Archaea

**DOI:** 10.1101/2022.04.06.487334

**Authors:** Logan H. Hodgskiss, Michael Melcher, Melina Kerou, Weiqiang Chen, Rafael I. Ponce-Toledo, Savvas N. Savvides, Stefanie Wienkoop, Markus Hartl, Christa Schleper

## Abstract

Ammonia oxidation as the first step of nitrification constitutes a critical process in the global nitrogen cycle. However, fundamental knowledge of its key enzyme, the copper-dependent ammonia monooxygenase is lacking, in particular for the environmentally abundant ammonia oxidizing archaea (AOA). Here, the structure of the enzyme is investigated by blue-native gel electrophoresis and proteomics from native membrane complexes of two AOA. Beside the known AmoABC subunits and the earlier predicted AmoX, two new protein subunits, AmoY and AmoZ, were identified. They are unique to AOA, highly conserved and co-regulated, and their genes are linked to other AMO subunit genes in streamlined AOA genomes. Modelling and in gel cross-link approaches support an overall protomer structure similar to the distantly related bacterial particulate methane monooxygenase indicating that AmoY and AmoZ serve an important structural and functional role. These data open avenues for further structure-function studies of this ecologically important key nitrification complex.

---

Nitrification, the conversion of ammonium to nitrate, is a crucial step in the global nitrogen cycle solely performed by microorganisms. The process has attracted particular attention due to its agricultural and environmental relevance. The first and rate limiting ^1^ step of nitrification is the oxidation of ammonia via the integral membrane protein complex ammonia monooxygenase (AMO) ^2,3^. While ammonia oxidizing bacteria (AOB) were first discovered over 125 years ago ^4^ and have been extensively studied, this biological process was also detected in the archaeal domain in the last 20 years ^5–7^. Ammonia oxidizing archaea (AOA) have gained broad attention as they are widespread in nature and are more abundant than their bacterial counterparts in most terrestrial and marine environments, indicating important roles in nitrogen cycling ^8–14^. Their central nitrogen and carbon metabolism, however, is distinct from that of AOB ^15–18^. In particular, subunits of the AMO complex show only about 40% identity to those of bacteria ^19^ and archaeal proteins catalyzing the second step in ammonia oxidation, i.e. the conversion of hydroxylamine to nitrite, are still unknown ^19–21^.

Due to the difficulty of growing nitrifying organisms and the inherent problems with isolating membrane proteins, no structural studies have been successfully carried out for any AMO complex, bacterial or archaeal. This holds true for most of the diverse enzymes of the CuMMO (copper-dependent membrane monooxygenase) protein family, with a few notable exceptions. Crystal structures ^22–26^ and one cryo-EM structure ^27^ of particulate methane monooxygenase (pMMO) from five methanotrophs have consistently confirmed a three-polypeptide protomer (subunits-A, -B and -C) arranged in a trimer of α_3_β_3_γ_3_ configuration with at least two conserved metal sites in each protomer. Even so, the elucidation of the active site has remained ambiguous. It was first proposed to reside in the PmoB subunit of pMMO ^28^ as recently supported by cryo-EM analysis ^27^, while differing amino acid conservation in Verrucomicrobia ^29^, a recent spectroscopic analysis ^30^, and mutagenesis of a hydrocarbon monooxygenase ^31^ suggest its localization in the PmoC subunit.

Although no AMO structure has been determined experimentally, homology modelling for the AMO of the bacterium *Nitrosomonas europaea* using pMMO as a template supported a homotrimeric structure as well as conservation of the Cu_B_ and Cu_C_ copper sites ^32^. The archaeal AMO complex is the most distantly related of all CuMMO proteins ^33,34^ and very little is known so far about its structure or function. Based on comparative metagenomics alone, it has been suggested that an additional subunit might be present in the complex, termed AmoX ^15,35^.

To gain insights into the overall architecture of the archaeal AMO complex, membrane protein fractions from the well characterized soil AOA, *Nitrososphaera viennensis*, were analyzed biochemically using native gel electrophoresis, mass spectrometry, and chemical cross linking. Beside the three known AmoABC proteins, three additional potential subunits were identified and one of the six predicted AmoC proteins in *N. viennensis* was recognized as the primary homolog in the protein complex. In addition, the overall subunit composition of the AMO complex was confirmed in the distantly related thermophilic AOA *Nitrosocaldus cavascurensis*.

## Results

### Complexome analysis of native membrane complexes displays the AMO composition of *Nitrososphaera viennensis*

*N. viennensis* was grown in continuous culture for several weeks under optimal growth conditions in order to obtain enough biomass for biochemical analyses (Melcher et al. in preparation). Between 800-2000 µg of membrane proteins were obtained from 450-550 mg of biomass per preparation, of which approximately 40-50 µg were loaded per lane on blue-native PAGE gels ^36^. After optimization of conditions, 22 bands were cut out and subjected to mass spectrometry (see Methods; Supplementary Fig. 1A). AMO subunits were among the most abundant proteins detected overall in these membrane fractions. The relative intensity profiles of AmoA, AmoB, and AmoC showed three distinct peaks corresponding to bands 4, 7, and 12, with the most prominent peak occurring at band 7 (Fig. 1A). The subunits AmoA, AmoB, and AmoC made up 10%, 5%, and 14%, respectively, of the total protein found in band 7 based on iBAQ normalized intensities. AmoX was also present in band 7 representing 10%. The most intense signals for the AmoC subunit were represented by two of the six AmoC homologs, AmoC6 and AmoC4. These two homologs could not be distinguished based on the peptides identified in the BN-PAGE gel. In denaturing SDS-Tricine-PAGE of cutouts from band 7 all known components of the AMO complex were visualized and confirmed by proteomics (Supplementary Fig. 2A). In addition, this allowed to identify unique peptides of the AmoC6 subunit (see Supplementary Discussion).

**Figure 1.**
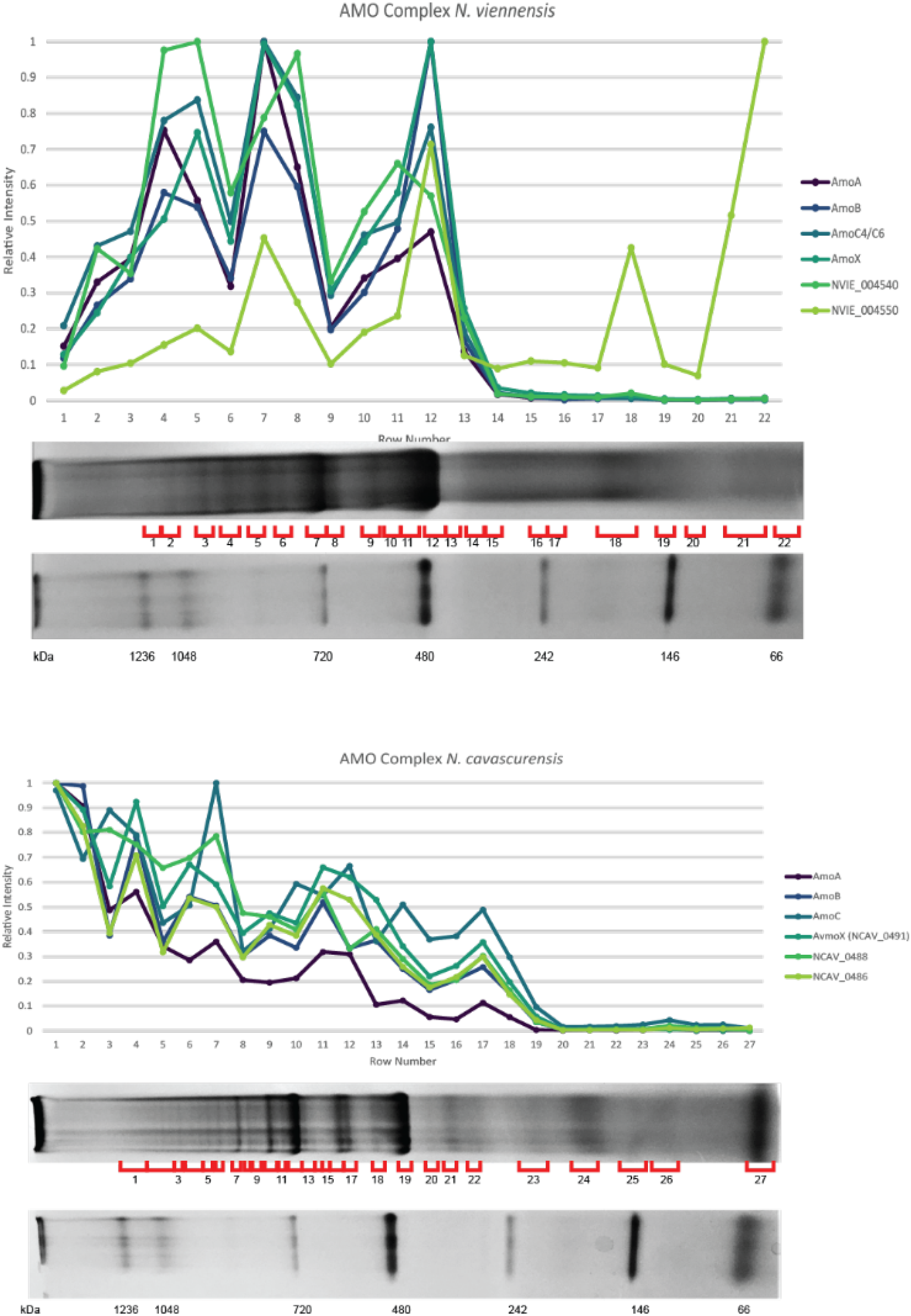
Relative intensity patterns of AMO subunits in BN-PAGE gels. Relative abundance of iBAQ normalized intensities of known and putative AMO subunits. iBAQ intensities for each protein are normalized to the highest detected intensity of that protein to create a relative abundance profile for each protein. **A.)** Patterns of AMO intensity in *N. viennensis*. **B.)** Patterns of AMO intensity in *N. cavascurensis*.

To identify additional proteins that might be part of the archaeal AMO complex a correlation analysis was conducted to find candidates with a similar migration pattern as all three primary AMO subunits AmoA, AmoB, and AmoC4/C6 in the BN-PAGE gel. Patterns of the 50% most abundant proteins were compared to each other using a Kendall correlation to determine the likelihood of dependence between various proteins. Additional criteria were (i) their presence in fully sequenced ammonia oxidizing archaea, and (ii) their absence in species that do not oxidize ammonia ^37^. Two proteins initially met these criteria: the putative AMO subunit AmoX and a hypothetical protein, NVIE_004540 (Table 1). The migration patterns for these proteins can be seen in Figure 1A. While this unbiased selection process produced intriguing additional AMO candidates, further analysis was needed to verify the presence of these newly identified and other potential subunits. Therefore, a multifaceted approach using genomics, proteomics, and transcriptomics was used to investigate this possibility.

**Table 1:** Correlations of proteins with occurrence of AmoA,B, and C in (A) *N. viennensis* and (B) *N. cavascurensis* BN-PAGE gels.

| Locus Tag | Gene | Protein* | Conserved in Extant AOA<sup>§</sup> | Exclusive to AOA<sup>§</sup> | Correlations<sup>†</sup> |  |  |
| --- | --- | --- | --- | --- | --- | --- | --- |
|  |  |  |  |  | AmoA | AmoB | AmoC6 |
| NVIE_016740 | NVIE_016740 | surface associated S-layer protein |  | X | X | X | X |
| NVIE_011620 | nuoJ | Complex I | X |  | X | X | X |
| NVIE_011600 | nuoM | Complex I | X |  | X | X | X |
| NVIE_027530 | coxB | Complex IV |  |  | X | X | X |
| NVIE_027540 | coxA1 | Complex IV |  |  | X |  |  |
| NVIE_013530 | NVIE_013530 | protein of unknown function |  | X | X | X | X |
| NVIE_027260 | NVIE_027260 | conserved protein of unknown function |  | X | X | X | X |
| NVIE_017130 | NVIE_017130 | protein of unknown function DUF373 | X |  | X |  | X |
| NVIE_027280 | amoX | potential AMO subunit | X | X | X | X | X |
| NVIE_004540 | NVIE_004540 | hypothetical protein | X | X | X | X | X |

B **AMO Correlation Results for *N. cavascurens***
| Locus Tag | Gene | Protein* | Conserved in Extant AOA<sup>§</sup> | Exclusive to AOA<sup>§</sup> | Correlations<sup>†</sup> |  |  |
| --- | --- | --- | --- | --- | --- | --- | --- |
|  |  |  |  |  | AmoA | AmoB | AmoC6 |
| NCAV_1585 | coxA | Cytochrome c oxidase polypeptide 1 |  |  | X | X |  |
| NCAV_1739 | NCAV_1739 | Uncharacterized protein |  | X | X | X |  |
| NCAV_1586 | coxB | Putative heme-copper oxidase subunit II |  |  | X | X | X |
| NCAV_0011 | NCAV_0011 | ABC-1 domain-containing protein |  |  | X | X |  |
| NCAV_1743 | amt | Ammonium transporter | X |  | X | X |  |
| NCAV_0191 | petB | Putative cytochrome b/b6 |  |  | X |  |  |
| NCAV_1587 | NCAV_1587 | Putative heme/copper-type cytochrome/quinol oxidase, subunit | X | X | X |  |  |
| NCAV_0486 | NCAV_0486 | Uncharacterized protein (NVIE_004550 homolog) | X | X | X | X | X |
| NCAV_0488 | NCAV_0488 | Uncharacterized protein (NVIE_004540 homolog) | X | X | X | X | X |
| NCAV_0491 | NCAV_0491 | Uncharacterized protein (AmoX) | X | X | X | X | X |

\*Proteins with at least one AmoABC protein correlation.
<sup>§</sup>Conservation and exclusiveness to AOA based on results from Abby et al. (2020).
<sup>†</sup>Correlation $\geq 0.7$ and adjusted p-value $\geq 0.001$ .

### Linkage analysis in AOA genomes supports proposed and additional AMO subunits

Earlier analyses of known subunits within the soil strains, or the family *Nitrososphaeraceae* (as defined by the Genome Taxonomy Database ^38^; used throughout), has shown a general lack of spatial clustering of all earlier known subunit genes. However, within the families *Nitrosopumilaceae* and *Nitrosocaldaceae*, the genes for the canonical AMO subunits, AmoABC, and the proposed subunit AmoX are syntenic ^35,39,40^. To investigate co-localization of potential additional subunit genes, the syntenic status and conservation across AOA of the 5 genes upstream and downstream of the AMO cluster in *Nitrosocaldaceae* and *Nitrosopumilaceae* were analyzed. Of these genes, 19 were conserved in AOA with 5 being found exclusively in AOA (Supplementary Data). The 5 genes of interest included two canonical AMO genes (*amo*A and *amo*B) and the genes *amo*X, NVIE_004540, and NVIE_004550. The *amo*X gene was previously identified in metagenomic studies ^15,35^ and NVIE_004540 was already a candidate identified from the BN-PAGE correlation analysis. The additional conserved protein, NVIE_004550, was newly identified and found to be located directly upstream of NVIE_004540, indicating potential co-transcription (Fig. 2). The two candidates encode for polypeptides of 9.6 kDa and 12.8 kDa respectively, and – like the candidate subunit AmoX - their predicted secondary structure is predominantly helical and their subcellular localization transmembrane.

**Figure 2.**
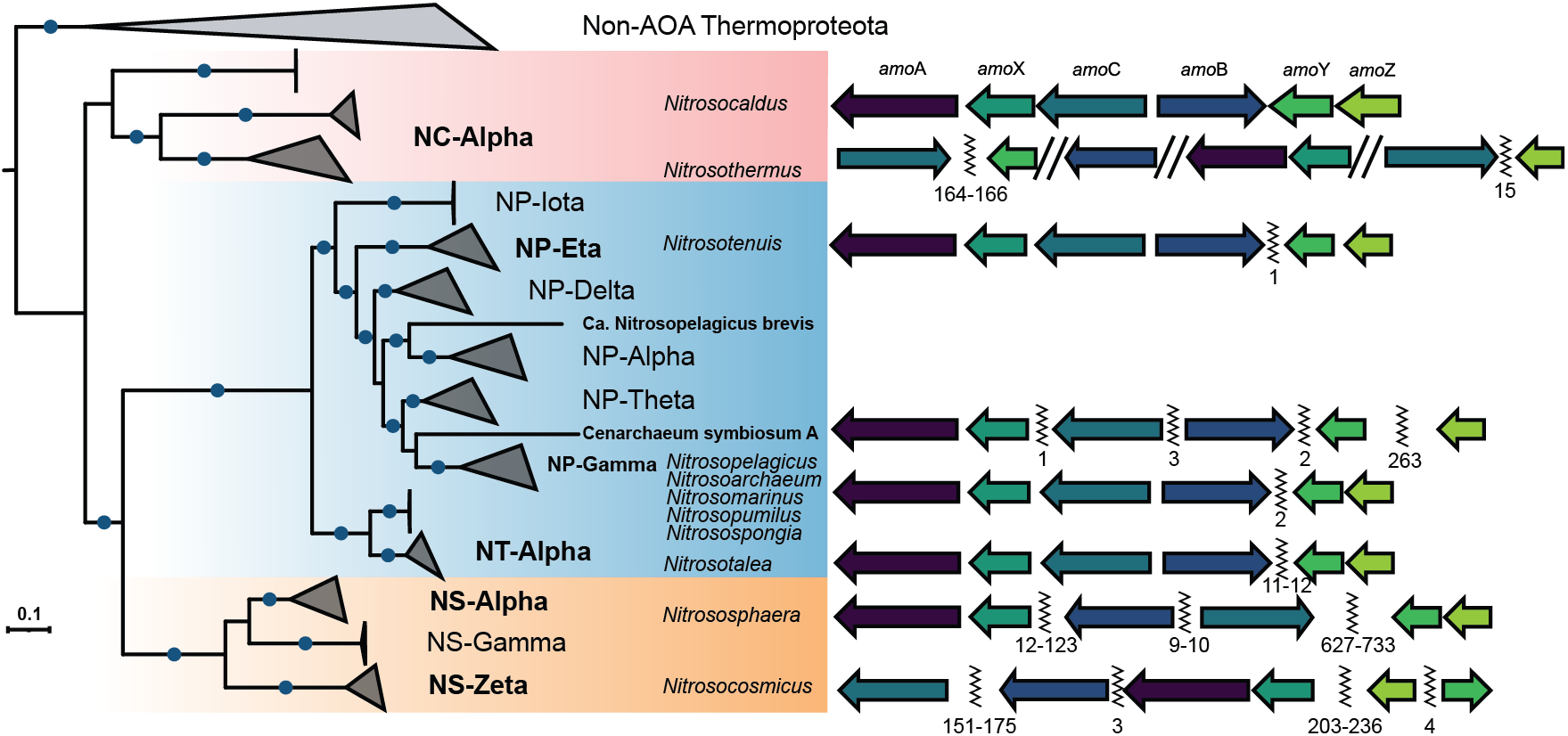
Genomic comparison of AMO subunit synteny in AOA. Left: phylogenetic tree of AOA based on 32 conserved ribosomal proteins, bootstrap values of 100% are indicated as blue circles. Taxonomic labels are colored according to GTDB family identity ^38^, *Nitrosocaldaceae*-red, *Nitrosopumilaceae*-blue, *Nitrososphaeraceae*-orange. Clades/organisms in bold were included in analysis. Clades are named according to Alves et al. (2018)^33^. Right: representation of general syntenic patterns in different clades of AOA. Homologs of NVIE_004540 are represented by AmoY and homologs of AmoZ are represented by AmoZ. Gaps between genes on the same contig are marked by a zig-zag line, double forward slash marks separate contigs. Numbers under the zig-zag lines represent number of genes between *amo* subunit genes. A full list of species and a finer analysis can be seen in Supplementary Fig. 10 and Supplementary Data.

A closer analysis in *Nitrosocaldaceae*, the earliest diverging lineage in evolutionary reconstructions of AOA ^37^, revealed that the genes for the three candidate subunits for AMO (AmoX, homolog of NVIE_004540, and homolog of NVIE_004550) clustered spatially with the canonical subunits (AmoABC) and were syntenic in *Nitrosocaldus cavascurensis* and *Ca*. Nitrosocaldus islandicus. Spatial clustering of all six subunit genes is also found in recently obtained MAGs ^41^ within the genus *Nitrosocaldus*. In the case of the newly proposed genus *Nitrosothermus* ^41^, AMO genes were split on multiple contigs and synteny could not be definitively determined (Fig. 2). Additionally, all six *amo* genes are predicted to have been newly acquired by the last common ancestor of AOA ^37^.

The emergence of *Nitrosopumilaceae* was accompanied by a separation of this genomic region into a primary cluster containing *amo*ABCX and a secondary cluster containing the homologs of NVIE_004540 and NVIE_004550 (Fig. 2). Within *Nitrosotalea* sp., these clusters are 11-12 genes apart, while the rest of *Nitrosopumilaceae* species have these clusters only 1-2 genes apart (with the exception of the sponge symbiont *Ca*. Cenarchaeum symbiosum). Apparently, the emergence of the family *Nitrososphaeraceae* led to a scattering of all subunit genes across the genome with the exception of *amo*A and *amo*X, which are typically linked.

A closer look was needed to account for the lack of association of NVIE_004550 with AmoABC in BN-PAGE of *N. viennensis*. When examining the relative abundance profile for NVIE_004550, the general pattern of AMO peptide peaks was followed. However, this remained undetected in the correlation analysis due to a high relative abundance peak occurring at the bottom of the gel peaking at the last band taken at approximately 66 kDa based on the BN-PAGE ladder (Fig. 1A). This is above the predicted mass of 12.8 kDa, but suggests that NVIE_004550 could also be part of the AMO but a possibly weaker association lead to its dissociation from the complex and migration to the bottom of the gel.

### BN-PAGE protein gel indicates same AMO composition in the thermophilic archaeon *Nitrosocaldus cavascurensis*

To test the composition of the AMO complex outside of the context of *N. viennensis*, the BN-PAGE approach was applied to membrane protein fractions of *N. cavascurensis*, a distantly related thermophilic AOA species of the *Nitrosocaldaceae* family ^39^ that was recently obtained in pure culture (Melcher et al. in preparation). Although a slightly different pattern of complexes was obtained (Fig. 1B) a correlation of the additional subunits was also observed with AmoA, AmoB, and AmoC in this thermophilic organism (Kendall correlation of proteins, as performed for *N. viennensis*). The three proteins AmoX, NCAV_0488 (homolog for NVIE_004540), and NCAV_0486 (homolog for NVIE_04550) all had migration patterns within the gel that strongly correlated with AmoABC (Table 1). This analysis confirmed that the proposed subunits were translated in *N. cavascurensis*, and potentially had a physical connection within the AMO complex.

### Chemical cross-linking supports physical interaction of additional subunits

To estimate the physical proximity of the proposed subunits to known subunits and other proteins within the BN-PAGE gel, in-gel cross-linking ^42^ was performed using the DSSO cross-linker on an additional BN-PAGE cut-out from band 7 (Supplementary Fig. 1B). Mass spectrometry and cross-linking analysis showed multiple cross-links among AmoA, AmoB, AmoC, and AmoX as well as with the two newly proposed subunits NVIE_004540 and NVIE_04550 (Fig. 3C). Many cross-links were also connected to NVIE_016740, a putative S-layer protein that likely represents a highly abundant surface layer protein as known from other archaea (SlaA) ^43,44^. As this protein presumably helps establish the pseudo-periplasm in AOA, it is not surprising to find it heavily cross-linked to membrane proteins.

**Figure 3.**
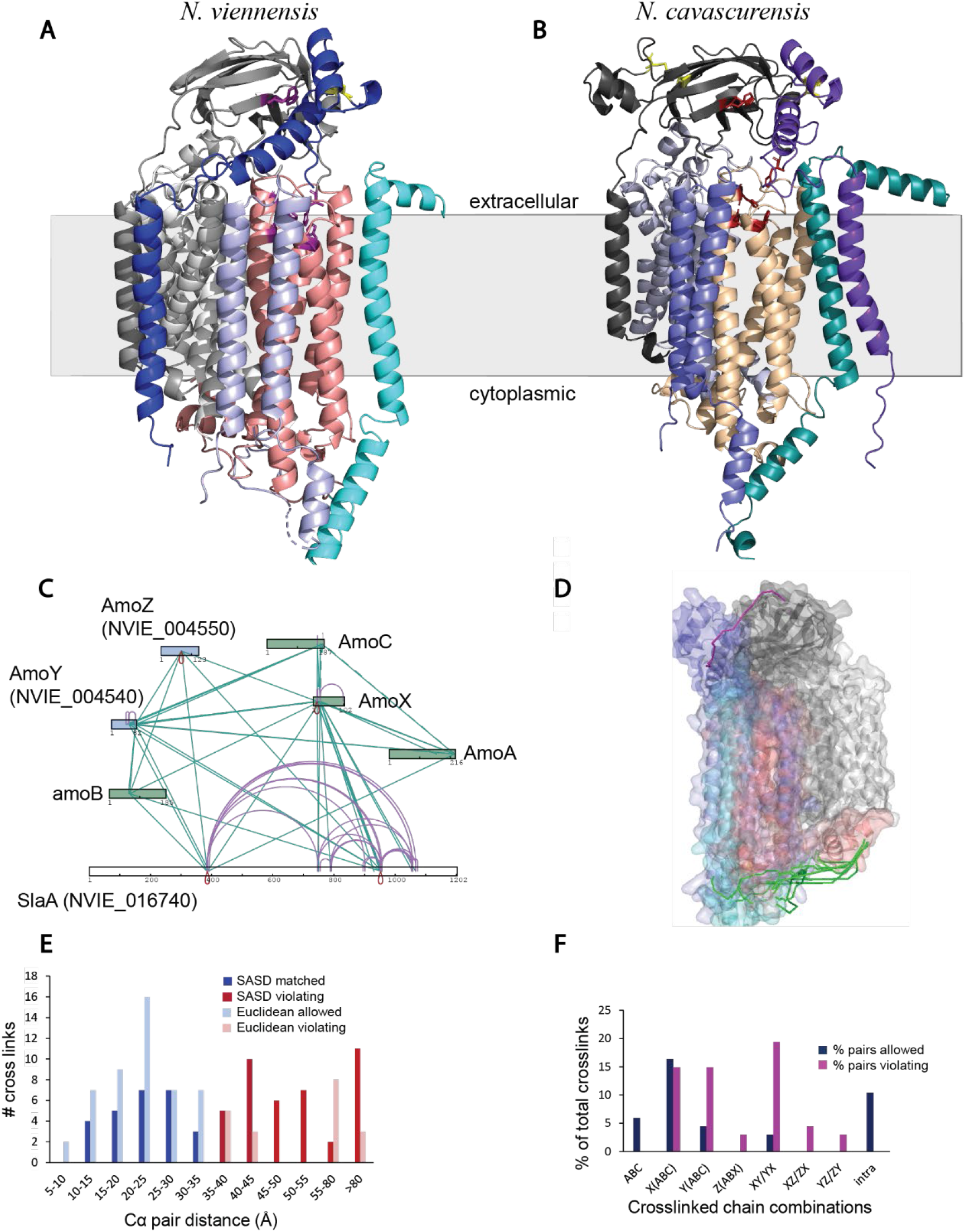
Structural support of proposed AMO subunits based on BN-PAGE cross-linking and Alpha Fold modelling. **A.)** Representation of identified cross-links among existing and proposed AMO subunits of an AMO band cut from a BN-PAGE gel of *N. viennensis*. Green: suspected subunits based on comparative genomics. Blue: newly proposed subunits based on BN-PAGE correlation and syntenic analysis. **B,C.)** Cartoon representations of the AlphaFold structure models of the NvAmoABCXYZ **(B)** and NcavAmoABCXYZ **(C)** hexamers, indicating their putative membrane orientation based on sequence hydropathy analysis. Subunits are colored as follows: NvAmoA, light grey; NvAmoB, grey; NvAmoC, salmon; NvAmoX, light blue; NvAmoY, cyan; NvAmoZ, blue; homologous subunits in Ncav are coloured in different shades of the same colour: NcAmoA, light grey; NcAmoB, dark grey; NcAmoC, sand; NcAmoX, sky blue; NcAmoY, teal; NcAmoZ, purple. Residues in the Cu_B_ and Cu_C_ copper sites are represented in magenta and red sticks in NvAMO and NcavAMO models respectively. Disulfide bonds are indicated in yellow. Note the different positioning of the AmoZ subunit. **D**.) Crosslinks within the SASD threshold for DSSO mapped on the NvAmoABCXY AlphaFold model depicted in green. The single observed crosslink between the AmoZ and AmoB subunits is depicted in magenta, as it violates distance criteria (50Å SASD) but is within range of Euclidean distance (31.8 Å). **E.)** Distribution of SASD and Euclidean Cα–Cα distances of unique DSSO crosslinks identified with Annika and MeroX. 27 out of 67 unique crosslinks satisfied distance criteria (SASD<35Å). **F.)** Percentage of crosslinked subunit combinations. While all crosslinks involving the canonical subunits AmoABC and approximately half of the crosslinks involving AmoX were satisfying distance criteria, only violating crosslinks were observed involving the putative subunit AmoZ.

AmoX also had individual cross-links to several other proteins (Supplementary Data). As only single connections were found, and these proteins did not appear in any other syntenic or correlative analyses, they were not taken to represent a structural role in the AMO complex. These cross-links can rather be attributed to the high abundance of those proteins in the cell membrane.

### Expression patterns of AMO subunits in *Nitrososphaera viennensis* and *Nitrosopumilus maritimus*

Available transcriptomic studies of AOA were inspected to explore whether the expression patterns of the newly predicted subunits would corroborate their involvement in the AMO. A recent study on copper limitation in *N. viennensis* ^45^ confirmed that the genes *amo*A, *amo*B, and *amo*C have some of the highest transcription levels in the cell, as also shown in previous studies ^46–48^. A clustering analysis of the same dataset revealed that *amo*A,B,C, *amo*X, NVIE_004540, and NVIE_004550 all appear to be co-regulated, and fell into the clusters containing the most highly expressed genes. (Supplementary Fig. 3, Supplementary Data).

A re-evaluation of these transcriptomic data (see Methods) also revealed *amo*C6 as the primarily transcribed *amo*C homolog (Fig. 4), thus confirming the identification of a unique AmoC6 peptide from an SDS band digested with chymotrypsin (Supplementary Discussion, Supplementary Data). Together this indicates that AmoC6 is the primary structural AmoC homolog in the AMO complex of *N. viennensis*, at least under the applied growth conditions.

**Figure 4.**
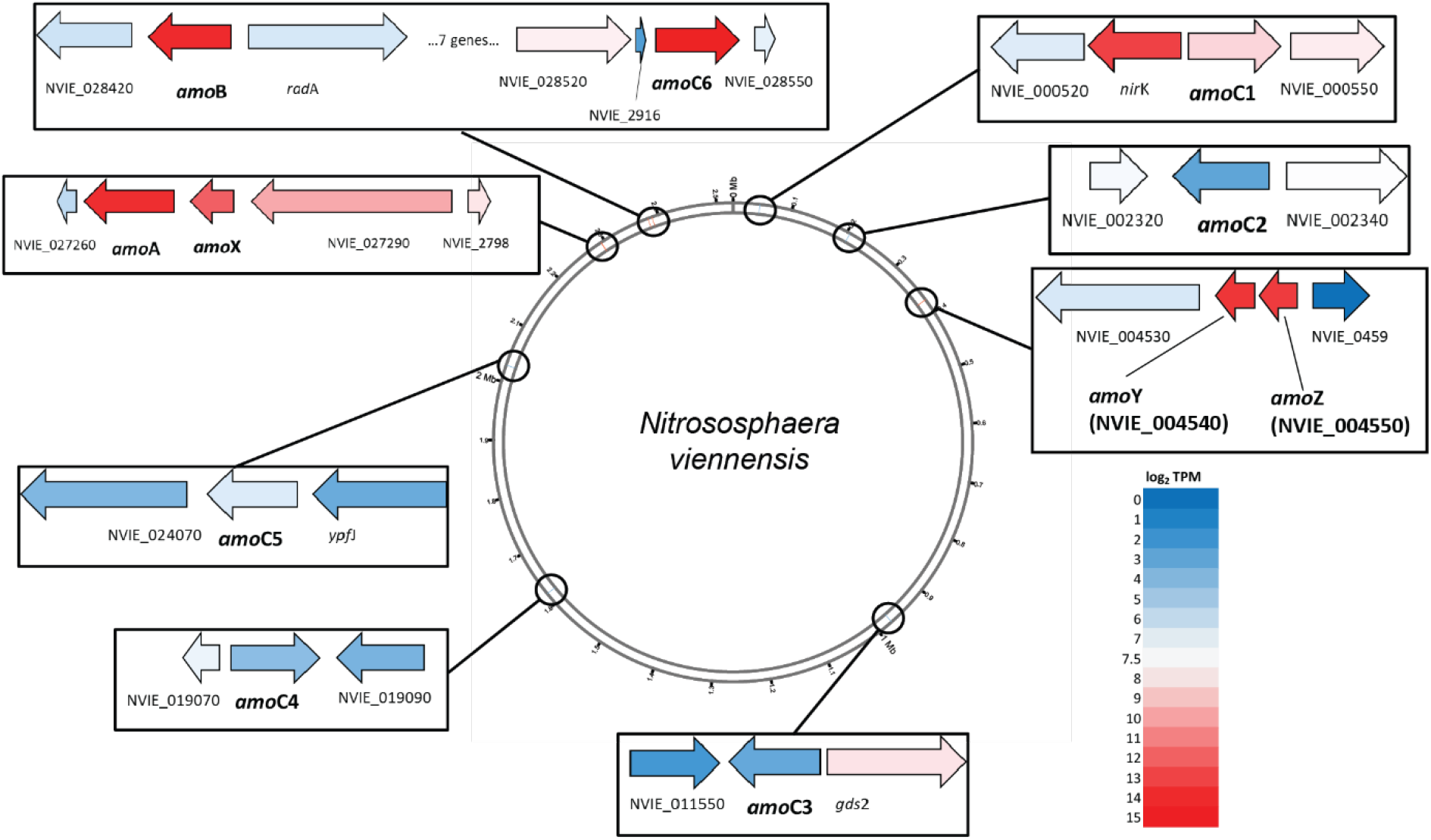
Transcription of AMO subunit genes in *N. viennensis*. Genomic representation of *N. viennensis* showing location of amo genes and average log_2_ transformed transcript per million(TPM) values from copper replete conditions in Reyes et al. (2020)^45^. Boxes show amo genes and immediate neighbors colored based on gene expression clusters from copper replete cultures. Red indicates a strong expression while blue represents a low or absent expression. All amo genes were found in clusters of highly expressed genes across both limited and replete conditions (see Supplementary Fig. 3).

Transcriptomics of the marine strain, *N. maritimus*, also showed high expression of *amo*A, *amo*B, *amo*C, *amo*X, and Nmar_1506 (homolog of NVIE_004540). Nmar_1507 (homolog of NVIE_004550), albeit syntenic with Nmar_1506, exhibited lower expression levels ^46^.

The three newly proposed AMO subunits were also inspected in proteomic datasets that were generated with methods allowing for the improved recovery of membrane proteins. All six of the known and proposed subunits were found in membrane fractions from *N. viennensis* from a previous study ^15^ as well as in the proteome of *N. maritimus* ^46^. In other proteomic studies of AOA ^49,50^, the three new subunits were not always present, likely due to their small size and limited number of trypsin cleavage sites.

### Structural search for missing components in the archaeal AMO complex

As previously observed, ^51^ comparisons of the amino acid sequences of the three subunits AmoA,B,C from archaea with those of bacteria indicate that the primary differences between the archaeal AMO subunits and the bacterial AMO subunits are missing transmembrane helices, at least one in AmoB and two in AmoC, and a C-terminal soluble portion found in AmoB/PmoB (Supplementary Figs. 4-6). These observations also hold true for the new clade of archaeal AMO recently discovered in the Thermoplasmata phylum ^52^. A HMMER search using the extended regions of the bacterial homologs against the genomes of collected AOA did not reveal any significant similarities. Therefore, a general structural search using Phobius ^53^ was carried out with the *N. viennensis* genome to search for genes that could encode a protein with the following criteria: (i) 1-3 transmembrane helices, (ii) conservation across all AOA ^37^, and (iii) present in the top 100 transcribed genes ^45^ (similar levels as the primary AMO subunits). This revealed six possible candidates (Table 2). The only candidates to meet the structural requirements while maintaining syntenic and similar patterns of migration in BN-PAGE were *amo*X, NVIE_004540, and NVIE_004550.

**Table 2:** Structural search for missing AMO subunits.

| Gene Expression & Structural Search |  |  |  | Previous Analyses |  | Protein |
| --- | --- | --- | --- | --- | --- | --- |
| Gene | TPM<br>log2* | TM<sup>†</sup> | AOA<br>Conservation<sup>§</sup> | BN-PAGE<br>Corr.<sup>‡</sup> | AMO<br>Synteny* |  |
| NVIE_013530 | 13.82 | 1 |  | X |  | protein of unknown function |
| NVIE_004540 | 13.55 | 1 | X | X | X | hypothetical protein |
| NVIE_004550 | 13.39 | 1 | X | X | X | hypothetical protein |
| amoX | 12.14 | 2 | X | X | X | putative ammonia monooxygenase - associated protein |
| coxB | 11.87 | 1 |  | X |  | putative heme-copper oxidase subunit II |
| NVIE_014370 | 11.75 | 1 |  |  |  | putative Copper binding protein, plastocyanin/azurin family |
| NVIE_010490 | 11.75 | 1 |  |  |  | putative phosphoesterase, DHHA1 |
| NVIE_027520 | 11.40 | 1 | X |  |  | putative heme/copper-type cytochrome/quinol oxidase, subunit |
| NVIE_027550 | 11.33 | 1 | X |  |  | putative blue (Type 1) copper domain protein |
| atpK | 11.30 | 2 |  |  |  | archaeal A1A0-type ATP synthase, subunit K |
| NVIE_021780 | 10.90 | 1 |  |  |  | exported protein of unknown function |
| NVIE_006540 | 10.90 | 1 |  |  |  | exported protein of unknown function |
| NVIE_029580 | 10.20 | 1 | X |  |  | blue (Type 1) copper domain-containing protein |

\*Based on transcriptomic counts averaged from replete copper conditions from Reyes et al. (2020).
<sup>†</sup>Number of predicted transmembrane helices.
<sup>§</sup>As predicted from Abby et al. (2020).
<sup>‡</sup>Represents proteins correlated with AmoA, AmoB, and AmoC in BN-PAGE bands from *N. viennensis* in this study.
\*Genes syntenic with *amoA*, *amoB*, and *amoC* based on the syntenic analysis from this study.

The addition of the three proposed subunits in archaea increases the number of transmembrane helices from 10-11 to approximately 14 per protomer making it comparable to the number found in bacterial crystal structures of pMMO where each protomer of the trimer (i.e. one unit of PmoABC), contains 14-15 transmembrane helices ^23,54^.

### Predicted structure of the archaeal AMO complex supports the integration of new subunits

To gain insights into the structural context of the archaeal AMO complex in the light of three additionally proposed subunits, a structural model for the organization of the *N. viennensis* AMO complex was obtained by employing the multimer-capable version of AlphaFold2.1 ^55–57^. The resultant models were all similar and represented confident predictions (top model, pLDDT=71.4 and ptm score=0.668). All predicted transmembrane helices from AmoX, NVIE_004540 (hereafter referred to as AmoY), and NVIE_04550 (hereafter referred to as AmoZ) play a role in anchoring the complex in the membrane along with the transmembrane helices from AmoA, AmoB, and AmoC (Fig. 3A). Additionally, the N-terminal end of AmoZ was predicted to contain two alpha helices that interact with the N-terminal domain of AmoB, thereby possibly replacing the role of the missing C-terminal soluble domain found in PmoB and offering the final piece of the missing complex in archaea (additional information in Supplementary Discussion). A disulfide bond was also predicted to form within the soluble domain of AmoZ. The overall structure is comparable to a protomer of the pMMO complex (Supplementary Fig. 7)

To compare the degree of conservation of the predicted hexameric organization of the AMO complex, a structural model of the AMO complex of *N. cavascurensis* was also obtained with AlphaFold2.1 (Fig. 3B). The resultant models were similar in their overall arrangement to each other and to the NvAmoABCXYZ model, with high overall confidence scores (top model, pLDDT=77.7 and ptm score=0.591). Notable differences between the *N*.*viennensis* and *N*.*cavascurensis* models include the localization of the transmembrane (TM) helix of AmoZ. In *N. viennenensis* the TM helix is predicted to interact mostly with the TM helix of AmoY, while in *N. cavascurenesis* it is predicted to interact with the TMs of AmoB and AmoA (Fig. 3A,B;Supplementary Fig. 8). This would affect the relative positioning of the N-terminal domain of AmoZ with respect to the AmoB soluble domain, allowing for a more “open” conformation. However, the extended loop connecting the N-terminal pair of helices in AmoZ with the TM domain theoretically allows for some flexibility (additional information in Supplementary Discussion).

Data from cross-linking experiments were mapped to the predicted model and strongly supported the predicted interactions (Fig. 3D) with some exceptions. Out of 67 unique observed cross-links, 27 (40%) satisfied a maximum solvent accessible surface distance (SASD) threshold of ≤ 35 Å (Fig. 3E), and involved all subunit combinations with the exception of AmoZ (Fig. 3F). AmoZ only participated in cross-linking interactions >35 Å, which supports a weaker association with the complex, as observed in the BN-PAGE migration patterns.

## Discussion

The archaeal AMO complex is a key enzyme of AOA energy metabolism that is highly expressed in all ammonia oxidizing organisms investigated and has large implications on the environment due to its overwhelming presence in many ecosystems ^8–14,58,59^. In the domain of Archaea, the only confirmed structural information stems from the crystal structure of a heterologously expressed AmoB originating from *Candidatus* Nitrosocaldus yellowstonensis ^60^. This structure confirmed the lack of the C-terminal cupredoxin domain and revealed an extended amino acid region not found in bacteria made up of two helices and two loops. It was proposed that this additional region could help stabilize the existing cupredoxin domain as supportive interactions are lacking due to the absence of the C-terminal domain. However, this amino acid extension is only found within the proposed genus of *Nitrosocaldus* (Supplementary Fig. 5). The work here profits from the recent improvements for cultivation of AOA in continuous cultures (Melcher et al. in preparation) and presents novel biochemical and comparative genomic evidence on the composition of the AMO complex in *Nitrososphaera viennensis* and other AOA.

The present analysis has verified that AmoX, NVIE_004540, and NVIE_004550 are all likely present within the archaeal AMO complex and proposes the naming of NVIE_004540 and NVIE_004550 as AmoY and AmoZ respectively. This finding is based on a host of analyses including proteomic, genomic, transcriptomic, structural, and modelling approaches.

In both *N. viennensis* and *N. cavascurensis*, the AMO complex migrated well above the predicted height of a homotrimeric complex, even when considering the additional subunits (predicted molecular weight of a homotrimeric complex with 6 subunits per protomer: 296.94 kDa *N. viennensis*; 305.101 kDa *N. cavascurensis*). This is in contrast to the PMO complex from a *Methylomirablis* species that was also extracted using n-dodecyl-β-D-maltoside (DDM) ^61^, and could be explained by their differences in membrane composition or potential differences in oligimerization of the protomer. AOA contain unique ether-linked lipids (i.e. crenarchaeols) ^62–67^ and rely on a proteinaceous S-layer rather than an outer membrane to create a pseudo-periplasmic space ^43,44^. The most likely explanation for the presence of three distinct peaks of AMO is the co-migration with other proteins or complexes that it could be physically interacting with, in particular with the S-layer protein.

Previous work on bacteria that rely on CuMMOs have identified other putative proteins involved with the complex. Notably, monocistronic transcripts containing *amo*ABC from the AOB *Nitrosococcus oceani* ATCC 19707 contained two additional genes assigned as *amo*R and *amo*D ^68^. *amo*R was found to be only present in *Nitrosococcus* and was therefore not thought to be a conserved part of bacterial AMO. A recent study indicated that *Amo*D/*Pmo*D (and the duplication *amo*E) play crucial roles in copper homeostasis, but they are not suspected to be a structural part of any CuMMO complex ^69^.

Although there is debate on whether AmoC harbors the primary active site in AMO, there is clear evidence that the metal site in PmoC plays a critical role in the complex of methanotrophs ^27,30,31^. While the archaeal AmoC lacks a substantial section found in all bacteria that corresponds to two transmembrane helices (Supplementary Fig. 6), the metal site is conserved across all archaeal and bacterial species and its importance is supported by site directed mutagenesis studies in the genetically tractable Actinobacteria that contain the homologous hydrocarbon monooxygenase ^31^.

These results are intriguing as the soil model AOA, *N. viennensis*, like most other soil dwelling AOA from the family *Nitrososphaeraceae*, encodes multiple homologs of the *amo*C gene while retaining only single copies of *amo*A and *amo*B ^15^ (Supplementary Data). Additional copies of *amo*C that are spatially disconnected from the AMO operon are encoded by some terrestrial AOB and were implicated in stress response based on transcriptional studies ^70,71^. Within *Nitrososphaeraceae*, no conserved AMO operons exist (Fig. 2). Duplications of the *amo*C gene (spatially distant from the other AMO genes) also occur in some species of the AOA marine associated family (*Nitrosopumilaceae)* and in two MAGs from AOA thermophiles (*Nitrosocaldaceae)*, all discovered in sediments ^41,72,73^. An *amo*C duplication is also found in an AOA sponge symbiont and copies of archaeal *amo*C are even found in marine viruses^74^. These findings together might indicate the metabolic importance of the AmoC subunit for ecophysiological adaptations in ammonia oxidation. While this work found AmoC6 to be the primary homolog within the complex for *N. viennensis*, it is possible that (some of) the other AmoC subunits, which arose by gene duplications at the species level (Supplementary Fig. 9), might be incorporated under certain environmental conditions and provide different activity profiles to the enzyme.

In conclusion, this study provides evidence through genomic, proteomic, and transcriptomic data for the presence of AmoX and the inclusion of AmoY and AmoZ as subunits within the archaeal AMO complex. A single protomer of the archaeal AMO would therefore consist of six subunits instead of three as in other complexes of the CuMMO family. As the anchoring of AMO in the membrane has previously been shown to be critical for its activity ^26^, it seems plausible that the newly identified subunits play an important role for the structural and functional integrity of AMO that allows it to properly function in archaea. The presence of a soluble domain within AmoZ that could replace the stabilizing function of the missing soluble domain in AmoB also fulfills a potentially crucial missing piece of the AMO complex. Definitive proof of the oligomerization and organization of these subunits will not be possible until a crystal structure of archaeal AMO is realized.

In the absence of additional structure-function analyses it remains an open question, in how far the additional subunits in the archaeal complex rather reflect the vast evolutionary distance to all other known protein complexes of the CuMMO family ^33^, or if this difference in structure also has relevant functional implications. For instance, the bacterial AMO complexes are promiscuous enzymes able to oxidize methane and other compounds ^75–78^. Such investigations on alternative substrates have not yet been performed with the archaeal complex, but would be important for evaluating the functional role of archaea in the environment.

Since AmoXYZ appear to have important structural roles it will be crucial to include all subunits in future expression and structural studies of this environmentally relevant protein complex. Considering the wide distribution of AOA in virtually all ecosystems ^8–14,33^ and their ecological relevance, developing genetic tools for AOA and improving their biomass production will be needed to enable structure-function analysis and to elucidate the full pathway of ammonia oxidation in these archaea.

## Methods

### AMO Alignments for amoABC

50 archaeal species’ and 29 bacterial species’ genomes were collected and searched for known AMO/PMO subunits (AmoA/PmoA, AmoB,/PmoB and AmoC/PmoC). Full lists of the collected species can be found in Supplementary Data.

If a species was not annotated, genomic.fna files were collected from NCBI and coding sequences were searched for using prodigal (version 2.6.3) ^79^ using the parameter -p single. In the case of annotated species, RefSeq annotation files were given preference if available. When no RefSeq annotation was available, GenBank annotations were used. In the case of *Ca*. Cenarchaeaum symbiosum A, the genome file was re-annotated using prodigal. This was done to search for coding regions that should theoretically be present that were not detected in the given annotation file.

Hidden Markov Models (HMMs) were made for archaea and bacteria separately based on amino acid sequences of well documented species with representatives from all major clades. In archaea, sequences from *Nitrososphaera viennensis* EN76 (*Nitrososphaeraceae*), *Nitrosocaldus cavascurensis* (*Nitrosocaldaceae*), *Nitrosopumilus maritimus* SCM1 (*Nitrosopumilaceae*, formerly Nitrosopumilales), and *Ca*. Nitrosotalea devaneterra (*Nitrosopumilaceae*, formerly Nitrosotaleales), were chosen to construct the model. In bacteria, sequences from *Nitrosococcus oceani* ATCC 19707 (γ-proteobacteria, ammonia oxidation), *Nitrosospira multiformis* ATCC 25196 (β-proteobacteria, ammonia oxidation), *Ca*. Nitrospira inopinata (Nitrospira, comammox), *Methylosinus trichosprium* OB3b (α-proteobacteria, methanotroph), *Methylococcus capsulatus* str. Bath (γ-proteobacteria, methanotroph), *Methylacidiphilum kamchatkense* Kam1 (Verrucomicrobia, methanotroph), and *Mycolicibacterium chubuense* NBB4 (Actinobacteria, hydrocarbon oxidation), were chosen to construct the model.

Sequences from representative species of archaea and bacteria were aligned using Mafft (version 7.427) ^80,81^ and an HMM model was constructed using hmmbuild (HMMER 3.3, hmmer.org) for each subunit in archaea and bacteria separately. An HMM search using hmmsearch (HMMER 3.3) was performed on selected species and sequences were collected for archaea and bacteria. Archaeal and bacterial species were searched separately due to the distant phylogentic relationship of the AMO/PMO complex within the two domains. A cut off value of 1e-20 was used for annotated species while a cut off value of 1e-10 was used for species analyzed with prodigal. A lower threshold was used for un-annotated species to account for the possibility of partial AMO/PMO genes on the edge of contigs. While a genome is not available for *Methylocystis* sp M., sequences were added to the appropriate bacteria files after the hmmsearch. Once collected, archaeal and bacterial sequences were combined and aligned using Mafft with the mafft-linsi paramter. Sequences that clearly did not belong after the multiple sequence (due to the presence of stop codons or inclusion due to the low threshold) were maunually removed (two in the case of bacterial AmoB/PmoB).

### *amo*C tree of archaeal genes

The archaeal *amo*C tree was constructed from a nucleotide BLAST (blastn, BLAST 2.12.0+) ^82^ search using sequences from AOA species representing the dominant clades of AOA (see above). Mafft was used to align the nucleotide sequences. The alignment was then trimmed using BMGE (v1.12) ^83^ and IQTree (version 2.1.2) ^84,85^ was used to construct the phylogeny of archaeal *amo*C using the ultra-fast method and a bootstrap value of 1000. Nucleotide sequences were used for the tree construction as amino acid sequences were too similar to construct a reliable phylogeny.

### Phylogenomic analysis

A total of 106 MAGs and completely sequenced genomes (98 AOA and 8 non-AOA genomes) were collected from NCBI, IMG or DDBJ databases, followed by protein prediction using Prodigal v2.6.3 ^79^. The identification of phylogenetic markers to perform the phylogenomic tree reconstruction was based on the workflow proposed by Graham et al. (2018) ^86^ using the archaeal single-copy gene collection (e-value 10^−10^) ^87^. 32 ribosomal proteins detected in at least 90 out of the 106 genomes present in the collected genome database were selected. Protein families were aligned independently using the mafft-linsi algorithm implemented in MAFFT v7.427 ^81^ followed by a trimming step in BMGE ^83^ with default parameters. Trimmed protein families were concatenated using a tailormade python script and the concatenated alignment was used to reconstruct a maximum likelihood (ML) phylogenomic tree in IQTREE (v2.0-rc1) ^84^ under the LG+C20+F+G model with 1000 bootstrap replicates.

### Reactor Growth

*N. viennensis* was grown as a continuous culture in 2 L bioreactors (Eppendorf) filled with 1.5 L of fresh water medium (FWM) ^65,88^ with modified trace element solution ^5^, 7.5 µM FeNaEDTA, 2 mM NH_4_Cl and 1 mM pyruvate at 42 °C and pH 7.5. Carbonate was supplied by gassing the reactors with a 98 % air 2 % CO_2_ mixture, and the applied dilution rates ranged from 0.035 to 0.07 h^-1^.

*N. cavascurensis* was grown as a batch culture in the same reactors, volume and medium as described for *N. viennensis*, but at 68 °C with 1 mM NH_4_Cl and pH 7.0. Carbonate was also supplied by gassing, but with a mixture of air/ N_2_/ CO_2_ to achieve a 10 % O_2_ and 2 % CO_2_ mixture. To increase the biomass, NH_4_Cl was added stepwise with syringes via a septum to increase the final NO_2_^-^ concentration to approximately 2.5 mM before harvesting the cultures.

Harvested biomass was concentrataed by centrifucation at 4°C and pellets were frozen at -70°C until further analysis.

### Membrane Protein Extraction

Procedures for protein extraction and running a BN-PAGE gel were based off of Witting et al. 2006 ^36^, Reisinger and Eichacker (2008) ^89^, and the NativePAGE™ Novex Bis-Tris Gel System manual from Life Technologies (MAN0000557). Study design and anlaysis for membrane extraction and BN-PAGE was also largely guided by de Almeida et al. (2016) ^90^ and Berger et al. (2021) ^91^.

Frozen pellets of biomass were thawed on ice and resuspended in a sodium phosphate buffer solution (50 mM sodium phosphate, 200 mM NaCl, pH 7.0) to a concentration of ∼ 20 mg/mL. Once resuspended, pepstatin and Complete Tablet EDTA-free inhibitor solution were added at concentrations of 1 µg/mL and 40 µL/mL (25x concentrated stock) respectively to inhibit protease activity. Cells were lysed using a One Shot machine set at 2.1 kbar of pressure. After lysis, samples were spun at 7000xg for 15 minutes at 4°C to remove cell debris. The supernatant was then taken for further processing.

Supernatant containing proteins and membrane were ultracentrifuged at 200,000xg (Beckman Coulter Ultracentrifuge; SW 41 Ti Swinging-Bucket Rotor, k_max_=124) for 90 minutes at 4°C using 13.2 mL thinwall polypropylene tubes with a level of deceleration set to 7. The supernatant (containing cytoplasmic proteins) was removed and stored at -70°C with 10% glycerol. The remaining membrane pellet was resuspended/washed in a solution of 1M NaCl, 50 mM Tris-HCl, pH 7.5 and then concentrated via ultracentrifugation at 200,000xg for 90 minutes at 4°C. The supernatant was again removed and stored at -70°C with 10% glycerol. A final washing of the membrane fraction was performed by resuspending the pellet in 50 mM Tris-HCl, pH 7.5 and then concentrated via ultracentrifugation at 200,000xg for 90 minutes at 4°C. (IMPORTANT: Washing solutions are made with Tris-Base and titrated with HCl to avoid accumulation of sodium ions that interfere with BN-PAGE gels. Do NOT make with Tris-HCl and titrate with NaOH.) After the final wash the supernatant was again removed and stored at -70°C with 10% glycerol.

The final membrane pellet was resuspended in 100-200 µL of NativePage Sample Buffer (Invitrogen BN2003) with 0.75 M of 6-aminocaproic acid. To aid in the resuspension of membrane fractions, samples were allowed to gnetly mix on a rotator set at 12 rpm at 4°C for 30 minutes. Following the incubation on the rotator, protein concentrations were measured using the Bradford assay (Bio-Rad #5000006). Based on the protein concentration, n-dodecyl-β-D-maltoside (DDM; Invitrogen BN2005) was added to the sample at a concentration of 0.5g DDM/g protein. Samples were again incubated on a rotator at 12 rpm and 4°C for 30 minutes. A final protein concentration was determined using the Bradford assay (Bio-Rad #5000006) with a bovine serum albumin (BSA) standard curve and controls to account for interference from DDM. Samples were then aliquoted into volumes containing approximately 40-50 µg of protein and frozen at -70°C for later analysis.

### BN-PAGE

Frozen membrane samples (40-50 µg aliquots; ∼10-20 µL) stored at -70°C were thawed on ice. Based on previously calculated protein concentrations, additional DDM was added to each sample to reach a DDM to protein ratio of 0.75g DDM / 1 g protein. Samples were then incubated in a shaker at 700 rpm for 15 minutes at 4°C. After incubation, samples were centrifuged at 9000xg for 60 minutes at 4°C to remove any cellular debris that was not solubilized by the addition of DDM. Supernatant from this centrifugation was transferred to a 1.5 mL LoBind protein tube. Coomassie (NativePage 5% G-250 Sample Additive; Invitrogen BN2004) was added to each sample to reach a Coomassie:DDM ratio of 1g:1g. Samples were then loaded on a 3-12% pre-cast BN-PAGE gel (Invitrogen BN1001). Approximately 5-7 µL of NativeMark Unstained Protein standard from Invitrogen (LC0725) was used as the ladder. The anode buffer was pre-ordered (Invitrogen BN2001) and consisted of a final concentration of 50 mM BisTris and 50 mM tricine at pH 6.8. The gel was run at 4°C in three stages. The first stage used dark blue cathode buffer (anode buffer with cathode buffer additive, Invitrogen BN2002; 50 mM BisTris, 50 mM tricine, 0.02% Coomassie G-250, pH 6.8) and was run for 1 hour at 150V. For the second stage, the dark blue cathode buffer was replaced with light blue cathode buffer (50 mM BisTris, 50 mM tricine, 0.002% Coomassie G-250, pH 6.8) and the gel was run for an additional hour at 250V. For the third and final stage, the light blue cathode buffer was replaced with anode buffer and run for 45 minutes at 250V.

When finished, gels were stained using SimplyBlue™ SafeStain (Invitrogen LC6060; maximum sensitivity protocol). Destaining was done with MilliQ (MilliporeSigma Milli-Q Reference A+ System) water and repeated until as much background could be removed as possible. Individual bands identified in each gel were removed and cut into 2-3 pieces and placed into LoBind protein Eppendorf tubes. Bands were stored at 4°C in 150 µL of MilliQ water until being processed for proteomic analysis, cross-linking, or SDS-Tricine-PAGE.

### SDS-Tricine-PAGE

Procedures for running an SDS-Tricine-Page gel were primarily based off of Schägger and Jagow (1987) ^92^ and Schägger (2006) ^93^. A 15% SDS-Tricine gel was made by mixing: 2.5 mL 30% acrylamide/Bis solution 37.5:1 (BioRad #1610158), 1.25 mL gel buffer (1.5 M Tris-HCl, 8.45 pH), 1.15 mL water, 50 μL 10% sodium-dodecyl sulfate (SDS), 50 μL 10% ammonium persulfate (APS), and 5 µL 1,2-bis(dimethylamino)ethan (TEMED). Once the gel solidified a 4% stacking gel consisting of 340 µL 30% acrylamide/Bis solution 37.5:1, 250 µL gel buffer (1.5 M Tris-HCl, 8.45 pH), 1.36 mL water, 20 μL 10% sodium-dodecyl sulfate (SDS), 20 μL 10% ammonium persulfate (APS), and 2 µL 1,2-bis(dimethylamino)ethan (TEMED) was poured on top. Selected BN-PAGE bands were cut into 2-3 pieces and incubated together in 20 µL SDS loading buffer (0.2 M Tris-HCl, 0.3 M dithiothreitol (DTT), 277 mM SDS (8% w/v), 6 mM bromophenol blue, 4.3 M glycerol) at 65°C for 90 minutes in a shaker at 500 rpm. The loading dye solution from this incubation was used to load the SDS-Tricine gel. Either 3-5 µL of Color Prestained Protein Standard-Broad Range (11-245 kDa) (New England Biolabs, P7712) or 1-3 µL of PageRuler Prestained Protein Ladder (10-180 kDa) (Thermo Scientific, 26617) were used as a ladder. Stock solutions of 10x concentrated anode buffer (1 M Tris-base, adjusted with 6M HCl to a pH of 8.9) and 10x concentrated cathode buffer (1 M Tris-base, 1 M Tricine, 1% SDS, pH 8.3 (no pH adjustment necessary)) were previously made. Buffers were diluted to 1x when used for running the gel. The gel was run at 30V for 25 minutes to allow the proteins to leave the stacking gel followed by 200V for ∼55 minutes or until the ladder reached the bottom of the gel. For clear visualization, SDS-Tricine gels were silver stained (see below). If the bands were to be used for proteomic analysis, gels were stained with SimplyBlue™ SafeStain (maximum sensitivity protocol). After staining, bands were cut from the gel and placed in LoBind protein epis with 150 µL of MilliQ water and stored at 4°C until being processed for proteomic analysis.

### Silver Staining of SDS-Tricine Gels

All steps were done using a gentle shaker. Gels were soaked in fixing solution (50% methanol, 12% acetic acid) for at least 1 hour to overnight. Containers with gels fixed overnight were sealed with parafilm to prevent evaporation. Gels were washed for 20 minutes in 50% ethanol. Washing was repeated twice for a total of 3 times. The gel was then soaked for 1 minute in a freshly prepared solution of 1.2 g/L of sodium thiosulfate pentahydrate. Next the gel was washed for 30 seconds in MilliQ water. This was repeated twice for a total of 3 times. After washing, the gel was soaked in the dark in freshly prepared silver staining solution (2 g/L silver nitrate, 0.04% formaldehyde) for 25-30 minutes. Following staining, the gel was washed twice with MilliQ water for 30 seconds. To develop bands, the gel was submerged in developer solution (60 g/L sodium carbonate, 0.04% formaldehyde, 0.036 g/L sodium thiosulfate pentahydrate). Bands developed within 1-3 minutes and development was stopped by adding destain solution (10% acetic acid, 1 % glycerol). The gel was then soaked in destain solution for approximately 5 minutes before being washed multiple times with MilliQ water.

### Silver Staining (Farmer’s Reducer) of SDS-Tricine Gels

The gel was soaked in fixing solution as stated above. Fixing solution was removed and the gel was soaked in Farmer’s Reducer (30 mM K_3_Fe(CN)_6_, 30 mM sodium thiosulfate pentahydrate) for 2 minutes. This will turn the gel yellow. The gel was then washed multiple times with MilliQ water until the yellow background was completely removed (30-90 minutes). Once the background was removed, water was poured off and replaced with 0.1% silver nitrate and was incubated in the dark for 15 minutes. After incubation, the gel was washed multiple times with MilliQ water. The gel was then submerged in a 2.5% sodium carbonate solution for 30 seconds. To develop bands, the solution was removed and replaced with a solution of 0.1% formaldehyde and 2.5% sodium carbonate. Development was stopped with destain solution as described above.

### DSSO Cross-Linking Blue Native PAGE cut-outs

The protocol from Hevler et al. was followed ^42^. To summarize, BN-PAGE cut-outs were immersed in 90 µL of sodium phosphate buffer solution (100 mM sodium phosphate, 0.15 M NaCl, pH 7.5: 42.6 mg NaH_2_PO_4_·H_2_O, 93.7 mg Na_2_HPO_4_, 0.3 mL 0.5M NaCl in 10 mL MilliQ water; pH adjusted to 7.5). 1 mg of the mass spec cleavable cross-linker disuccinimidyl sulfoxide (DSSO;Thermo Scientific A33545) was resuspended in 51.5 µL of dimethyl sulfoxide (DMSO) to reach a final concentration of 50 mM DSSO. 10 µL of 50 mM DSSO was then added to each sample yielding a final DSSO concentration of 5 mM per sample. Samples were briefly vortexed and then incubated at room temperature for 30 minutes. The cross-linking reaction was then stopped with the addition of 2 µL of 1 M Tris-HCl. Samples were briefly vortexed and incubated for 15 minutes at room temperature. The solution was then removed and replaced with MilliQ water. Samples were then stored at 4°C until being processed for mass spectrometry analysis. While results are convincing, in gel chemical cross-linking is a relatively new technique ^42^, and putative artifacts cannot be excluded due to the lack of a strong control.

### Sample preparation for mass spectrometry (SDS or BN-gel)

The Coomassie-stained gel bands were destained with a mixture of acetonitrile (Chromasolv®, Sigma-Aldrich) and 50 mM ammonium bicarbonate (Sigma-Aldrich). The proteins were reduced using 10 mM dithiothreitol (Roche) and alkylated with 50 mM iodoacetamide. Trypsin (Promega; Trypsin Gold, Mass Spectrometry Grade) digestion was carried out at 37°C overnight in 50mM ammonium bicarbonate. Chymotrypsin (Roche) digestion was carried out at 25°C for 5 hours in 50mM ammonium bicarbonate. GluC (Roche) digestion was carried out at 37°C overnight in 50mM ammonium bicarbonate. 10% formic acid was used to stop the digestion and peptides were extracted twice with 5% FA for 10min in a cooled ultrasonic bath. Extracted peptides were pooled and desalted using C18 Stagetips ^94^.

### Liquid chromatography separation coupled to mass spectrometry

Peptides were analyzed on an UltiMate 3000 HPLC RSLC nanosystem (Thermo Fisher Scientific) coupled to a Q Exactive HF-X, equipped with a nano-spray ion source using coated emitter tips (PepSep, MSWil). Samples were loaded on a trap column (Thermo Fisher Scientific, PepMap C18, 5 mm × 300 μm ID, 5 μm particles, 100 Å pore size) at a flow rate of 25 μL min^-1^ using 0.1% TFA as the mobile phase. After 10 min, the trap column was switched in-line with the analytical C18 column (Thermo Fisher Scientific, PepMap C18, 500 mm × 75 μm ID, 2 μm, 100 Å) and peptides were eluted by applying a segmented linear gradient from 2% to 80% solvent B (80% acetonitrile, 0.1% formic acid; solvent A 0.1% formic acid) at a flow rate of 230 nL/min over 60 min. The mass spectrometer was operated in data-dependent mode, survey scans were obtained in a mass range of 350-1600 m/z with lock mass activated, at a resolution of 120,000 at 200 m/z and an AGC target value of 1E6. The 15 most intense ions were selected with an isolation width of 1.2 Thomson for a max. of 150 ms, fragmented in the HCD cell at stepped normalized collision energy at 26%, 28%, and 30%. The spectra were recorded at an AGC target value of 1E5 and a resolution of 60,000. Peptides with a charge of +1, or >+7 were excluded from fragmentation, the peptide match feature was set to preferred, the exclude isotope feature was enabled, and selected precursors were dynamically excluded from repeated sampling for 20 seconds within a mass tolerance of 8 ppm.

### Data analysis for identification of BN-PAGE and SDS-Tricine-PAGE bands

For peptide and protein identification raw data were processed using the MaxQuant software package ^95^ (version 1.6.6.0) and spectra searched against Nitrososphaera_viennensis reference proteome (Uniprot, downloaded fall 2021) with a starting site modification (see below) to AmoC4 (Uniprot accession: A0A060HLS1) and a database containing common contaminants. The search was performed with full trypsin specificity (or corresponding enzyme used in the digestion) and a maximum of 2 missed cleavages at a protein and peptide spectrum match false discovery rate of 1%. Carbamidomethylation of cysteine residues was set as a fixed modification and oxidation of methionine and N-terminal acetylation as variable modifications. The output option of iBAQ with log fit was selected. All other parameters were left at default.

### Data analysis for identification of BN-PAGE cross-linked bands

Peptide and protein identification was performed as described as except with MaxQuant version 1.6.17.0 and with the a reference proteome that was not corrected for AmoC4.

To identify cross-linked peptides, the raw data were searched with either MS Annika ^96^ in Proteome Discoverer 2.3 or with MeroX 2.0 ^97^ against the sequences of the top abundant protein hits (with at least 10 MS/MS counts) from the MaxQuant search. Although it had less than 10 MS/MS counts, the protein encoded by NVIE_004550 was also added based on other proteomic and syntenic analysis. DSSO was selected as the cross-linking chemistry. Carbamidomethyl on Cys was set as a fixed modification and oxidation of Met and protein N-terminal acetylation as variable modifications. Enzyme specificity was selected according to the protease used for digestion. Search results were filtered for 1% FDR on the PSM level limiting the precursor mass deviation to 10 ppm. Further filtering was done for only non-decoy and high confidence PSMs in MS Annika and for a score higher than 50 in MeroX 2.0. http://crosslinkviewer.org/ was used to draw the XL maps.

Solvent Accessible Surface Distances (SASD) between crosslinked residues were calculated and plotted on the AlphaFold *Nv*AmoABCXYZ structure model with Jwalk, and scored with the MNXL program to assess whether they violate distance criteria ^98^. Crosslinks were considered “matched” if the SASD between the crosslinked residues was <35Å (based on Cα-Cα distances).

### AmoC4 start site in *N. viennensis*

Six homologs of *amo*C (*amo*C1-6) are present in the genome of *N. viennensis*. Predicted start sites for AmoC1-3, AmoC5, and AmoC6 are consistent across annotations (GenBank and RefSeq; Supplementary Fig. 2). However, the start site of AmoC4 is significantly different between the two annotations. Additionally, there is a third possibility based on a methionine that would resemble the start site of AmoC1-3,5-6. Based on AmoC alignments in all AOA and transcriptional information from Reyes et al. (2020), it was determined that the most accurate start site for this protein resides at the third option, resembling that of the other AmoC subunits in *N. viennensis*. The proteome downloaded from Uniprot was manually annotated to reflect this decision. This is the proteome used for BN-PAGE and SDS-Tricine-PAGE reference files (not for cross-linked analysis). With this correction, it is not possible to distinguish between AmoC4 and AmoC6 in any of the samples. However, if the original proteome from Uniprot is used, MaxQuant will identify unique peptides for AmoC4 and AmoC6. Based on the current analysis, this is an inaccurate interpretation of the data. With the corrected AmoC4 annotation, only unique peptides from AmoC6 are found. To further verify that AmoC4 is not playing a significant role, the raw data from the chymotrypsin digest was subjected to an unspecific and semi-specific closed search for unique peptides from AmoC4. An open search was also carried out using the FragPipe software (version 17.1) ^99^. None of these searches revealed unique AmoC4 peptides therefore strengthening the argument that the primary structural homolog in *N. viennensis* is AmoC6. Peptide coverage maps for the three digests (trypsin, GluC, and chymotrypsin) were created using Protein Coverage Summarizer (v1.3.8056) (https://github.com/PNNL-Comp-Mass-Spec/protein-coverage-summarizer/releases).

### BN-PAGE correlation analysis

A Kendall correlation using R was used to find proteins that had a similar migration pattern with AmoABC. Any protein expected to be a part of the AMO complex should be in high abundance. Therefore, only the top 50% proteins, based on iBAQ abundance, were used for the analysis. Each protein was correlated with every other protein using the function cor.test in R with method=“kendall” and use=“complete.obs” using iBAQ values. P-values were adjusted using the Benjamini-Hochberg method. Results were filtered for proteins correlated with one of the AMO proteins (AmoA, AmoB, or AmoC) and a correlation (tau) greater than or equal to 0.7. All filtered results had an adjusted p-value higher than 0.001.

For genes of interest, conservation across AOA was checked according to the data set curated in Abby et al. (2020)^37^. A gene was considered exclusive to AOA if it was found in AOA but not found in any species outside of AOA according to the dataset in Abby et al. (2020)^37^. The full dataset from this study was provided by the authors and can be found with other datasets here (link provided upon publication).

### Transcriptomic Clustering

Transcriptomic reads from Reyes et al. (2020) ^45^ were re-processed taking into account strandedness and using hisat2 (version 2.1.0) ^100^ rather than bowtie2 for read mapping. Count values were evaluated using featureCounts (v2.0.0) ^101^. Counts were then normalized using the Transcripts per Kilobase Million (TPM) method. TPM is calculated by dividing each gene count by the total length of the gene in kilobases giving a reads per kilobase value. “reads per kilobase” was summed up for all genes in a sample and divided by 1,000,000 to give a scaling factor. Each genes’ “reads per kilobase” was divided by this scaling factor to give the final TPM. TPM values were converted to log_2_ values and clustered using hierarchal clustering with the default values of heatmap.2 ^102^ in R version 3.6.3 ^103^. Clustered genes were split into 15 clusters. 3 clusters represented the genes with the highest abundance of transcripts in both the copper replete and copper limited cultures. Cluster data is visualized using Anvio 7.1 ^104^ (Supplementary Fig. 3).

### Structural Search for Missing AMO Sections in Archaea

Extended pieces of bacterial AmoB and AmoC were trimmed from alignments of combined archaea and bacteria species using trimal (v1.4rev15) ^105^. One sequence was removed from AmoB sequences as it was not actually the AmoB subunit. A second was removed due to the inclusion of stop codons in the sequence. Trimmed sections were used to create HMMs and then searched against archaeal species to look for proteins that might supply these missing pieces. No hits were found. Therefore, a structural search for proteins containing transmembrane helices in *N. viennensis* was carried out for highly transcribed genes.

TPM reads from Reyes et al. 2020^45^ were averaged for the five replete conditions and sorted by highest abundance to obtain the top 100 transcribed genes. The GenBank translated CDS region for *N. vienneneis* was analyzed using Phobius ^53^ to identify transmembrane helices and signal peptides in all proteins. The highest transcribed genes were then filtered to include genes with 1-3 predicted transmembrane helices. Candidate genes were then analyzed for conservation in AOA and correlation in BN-PAGE gels from *N. viennensis*.

### Predicted model of archaeal AMO using AlphaFold-multimer

Sequences for AmoA, AmoB, AmoC, AmoX, NVIE_004540 (AmoY), and NVIE_004550 (AmoZ) from *N. viennensis* were used for AlphaFold2.1 ^55–57^ predictions. In the case of AmoB and NVIE_004550, predicted signal peptides were removed based on predictions using SignalP 5.0 (archaea) ^106^. AmoC6 was used to represent the AmoC subunit. For *N. cavascurensis*, sequences for AmoA, AmoB, AmoC, NCAV_0491 (AmoX), NCAV_0488 (homolog of NVIE_004540; AmoY), and NCAV_0486 (homolog of NVIE_004550; AmoZ) were used after predicted signal peptides were removed based on predictions using SignalP 5.0 (archaea) ^106^. All images were generated in PyMOL ^107^.

### Identification and Alignments of amoXYZ in Ammonia Oxidizing Archaea

Amino acid sequences for AmoX, AmoY, and AmoZ were obtained from the genes *amo*X, NVIE_004540, and NVIE_004550 respectively. Homologs in other AOA were initially searched for using blastp (v2.12.0+) ^82^ with a threshold of 1e-4. Not all species had identified hits. A more sensitive analysis was performed by creating an HMM model using hmmbuild (HMMER 3.3, hmmer.org) for each new subunit from the top BLASTp hit from each species with a BLAST result. The HMMs for AmoX, AmoY, and AmoZ were then used with hmmsearch (HMMER 3.3, hmmer.org) in all collected AOA genomes. This produced hits in all species for all subunits except amoY and amoZ in Thaumarchaeota archaeon J079, a MAG that is also missing AmoB and is only 84% complete, and amoZ in the GenBank protein file for *Ca*. Cenarchaeum symbiosum A. A re-analysis of the *Ca*. C. symbiosum A genome using prodigal was able to identify a coding sequence for AmoZ while maintaining the coding sequences for all other AMO subunits. A complete list of all identified AMO subunits in the collected species can be found here (link provided upon publication).

## Data and Code Availability

All proteomic data was deposited to the ProteomeXchange Consortium via PRIDE ^108^ partner repository (number provided upon publication). Relevant scripts and code for data analysis can be found at (link provided upon publication).

## Supplementary Information

### Supplementary Discussion

Discussion on SDS-Tricine-Page results and differences between *N. viennensis* and *N. cavascurensis* AlphaFold2.1 models.

**Supplementary Data 1** (available upon publication)

Proteomic data and supplementary tables.

**Supplementary Data 2** (available upon publication)

Genomic and transcriptomic data and supplementary tables.

**Source Data 1** (available upon publication)

A .pdb file for the predicted AlphaFold2.1 structure of *N. viennensis* (Alphafold_NvAMO_rank_1.pdb).

**Source Data 2** (available upon publication)

A .pdb file for the predicted AlphaFold2.1 structure of *N. cavascurensis* (NcavAMO_rank_1.pdb).

## Acknowledgements

We thank Anas Mohammed Mardini for excellent technical assistance in the cultivation of N. viennensis and Wolfram Weckwerth for valuable input in the initial discussions of the project. We also thank Florian Sikora and Dr. Boris Görke for technical assistance and usage of the OneShot machine for cell lysis and Dr. Stephanie Eichorst for assistance and usage of the ultracentrifuge. We are also appreciative of Dr. Thomas Rattei, Florian Goldenberg, and Johann Dorn of the Division of Computational Systems Biology (CUBE) for providing maintenance and access to the Life Science Computer Cluster (LiSC) at the University of Vienna. This project was supported by Doktoratskolleg (DK) plus: Microbial nitrogen cycling – from single cells to ecosystems (Austrian Science Fund W1257) and by ERC Advanced Grant TACKLE (No. 695192).

## References

1. Wong-Chong, G. M. & Loehr, R. C. The kinetics of microbial nitrification. Water Res. 9, 1099–1106 (1975).

2. Hyman, M. R. & Wood, P. M. Suicidal inactivation and labelling of ammonia mono-oxygenase by acetylene. Biochem. J. 227, 719–725 (1985).

3. Hollocher, T. C., Tate, M. E. & Nicholas, D. J. Oxidation of ammonia by Nitrosomonas europaea. Definite 18O-tracer evidence that hydroxylamine formation involves a monooxygenase. J. Biol. Chem. 256, 10834–10836 (1981).

4. Winogradsky, S. Recherches sur les organismes de la nitrification. Ann Inst Pateur 4:213–231., (1890).

5. Könneke, M. et al. Isolation of an autotrophic ammonia-oxidizing marine archaeon. Nature 437, 543–546 (2005).

6. Treusch, A. H. et al. Novel genes for nitrite reductase and Amo-related proteins indicate a role of uncultivated mesophilic crenarchaeota in nitrogen cycling. Environ. Microbiol. 7, 1985–1995 (2005).

7. Venter, J. C. et al. Environmental Genome Shotgun Sequencing of the Sargasso Sea. Science (80-.). 304, 66–74 (2004).

8. Leininger, S. et al. Archaea predominate among ammonia-oxidizing prokaryotes in soils. Nature 442, 806–809 (2006).

9. Nicol, G. W., Leininger, S., Schleper, C. & Prosser, J. I. The influence of soil pH on the diversity, abundance and transcriptional activity of ammonia oxidizing archaea and bacteria. Environ. Microbiol. 10, 2966–2978 (2008).

10. Adair, K. L. & Schwartz, E. Evidence that ammonia-oxidizing archaea are more abundant than ammonia-oxidizing bacteria in semiarid soils of northern Arizona, USA. Microb. Ecol. 56, 420–426 (2008).

11. Karner, M. B., Delong, E. F. & Karl, D. M. Archaeal dominance in the mesopelagic zone of the Pacific Ocean. Nature 409, 507–510 (2001).

12. Shi, Y., Tyson, G. W., Eppley, J. M. & Delong, E. F. Integrated metatranscriptomic and metagenomic analyses of stratified microbial assemblages in the open ocean. ISME J. 5, 999–1013 (2011).

13. Baker, B. J., Lesniewski, R. A. & Dick, G. J. Genome-enabled transcriptomics reveals archaeal populations that drive nitrification in a deep-sea hydrothermal plume. ISME J. 6, 2269–2279 (2012).

14. Hollibaugh, J. T., Gifford, S., Sharma, S., Bano, N. & Moran, M. A. Metatranscriptomic analysis of ammonia-oxidizing organisms in an estuarine bacterioplankton assemblage. ISME J. 5, 866–878 (2011).

15. Kerou, M. et al. Proteomics and comparative genomics of Nitrososphaera viennensis reveal the core genome and adaptations of archaeal ammonia oxidizers. Proc. Natl. Acad. Sci. 113, E7937–E7946 (2016).

16. Walker, C. B. et al. Nitrosopumilus maritimus genome reveals unique mechanisms for nitrification and autotrophy in globally distributed marine crenarchaea. Proc. Natl. Acad. Sci. 107, (2010).

17. Berg, I. A., Kockelkorn, D., Buckel, W. & Fuchs, G. A 3-Hydroxypropionate/4-Hydroxybutyrate Autotrophic Carbon Dioxide Assimilation Pathway in Archaea. Science (80-.). 318, (2007).

18. Könneke, M. et al. Ammonia-oxidizing archaea use the most energy-efficient aerobic pathway for CO2 fixation. Proc. Natl. Acad. Sci. U. S. A. 111, 8239–44 (2014).

19. Lancaster, K. M., Caranto, J. D., Majer, S. H. & Smith, M. A. Alternative Bioenergy: Updates to and Challenges in Nitrification Metalloenzymology. Joule (2018) doi:10.1016/j.joule.2018.01.018.

20. Simon, J. & Klotz, M. G. Diversity and evolution of bioenergetic systems involved in microbial nitrogen compound transformations. Biochim. Biophys. Acta -Bioenerg. 1827, 114–135 (2013).

21. Kozlowski, J. A., Stieglmeier, M., Schleper, C., Klotz, M. G. & Stein, L. Y. Pathways and key intermediates required for obligate aerobic ammonia-dependent chemolithotrophy in bacteria and Thaumarchaeota. ISME J. 1–10 (2016) doi:10.1038/ismej.2016.2.

22. Lieberman, R. L. & Rosenzweig, A. C. Crystal structure of a membrane-bound metalloenzyme that catalyses the biological oxidation of methane. Nature 434, 177– 182 (2005).

23. Hakemian, A. S. et al. The metal centres of particulate methane mono-oxygenase from Methylosinus trichosporium OB3b. Biochem. Soc. Trans. 36, 1134–1137 (2008).

24. Smith, S. M. et al. Crystal structure and characterization of particulate methane monooxygenase from Methylocystis species strain M. Biochemistry 50, 10231–10240 (2011).

25. Sirajuddin, S. et al. Effects of zinc on particulate methane monooxygenase activity and structure. J. Biol. Chem. 289, 21782–21794 (2014).

26. Ro, S. Y. et al. From micelles to bicelles: Effect of the membrane on particulate methane monooxygenase activity. J. Biol. Chem. 293, 10457–10465 (2018).

27. Chang, W. H. et al. Copper Centers in the Cryo-EM Structure of Particulate Methane Monooxygenase Reveal the Catalytic Machinery of Methane Oxidation. J. Am. Chem. Soc. 143, 9922–9932 (2021).

28. Balasubramanian, R. et al. Oxidation of methane by a biological dicopper centre. Nature 465, 115–119 (2010).

29. Op den Camp, H. J. M. et al. Environmental, genomic and taxonomic perspectives on methanotrophic Verrucomicrobia. Environ. Microbiol. Rep. 1, 293–306 (2009).

30. Ross, M. O. et al. Particulate methane monooxygenase contains only mononuclear copper centers. Science (80-.). 364, 566–570 (2019).

31. Liew, E. F., Tong, D., Coleman, N. V. & Holmes, A. J. Mutagenesis of the hydrocarbon monooxygenase indicates a metal centre in subunit-C, and not subunit-B, is essential for copper-containing membrane monooxygenase activity. Microbiol. (United Kingdom) 160, 1267–1277 (2014).

32. Musiani, F., Broll, V., Evangelisti, E. & Ciurli, S. The model structure of the copper-dependent ammonia monooxygenase. JBIC J. Biol. Inorg. Chem. 1, 3 (2020).

33. Alves, R. J. E., Minh, B. Q., Urich, T., Von Haeseler, A. & Schleper, C. Unifying the global phylogeny and environmental distribution of ammonia-oxidising archaea based on amoA genes. (2018) doi:10.1038/s41467-018-03861-1.

34. Khadka, R. et al. Evolutionary History of Copper Membrane Monooxygenases. (2018) doi:10.3389/fmicb.2018.02493.

35. Bartossek, R., Spang, A., Weidler, G., Lanzen, A. & Schleper, C. Metagenomic Analysis of Ammonia-Oxidizing Archaea Affiliated with the Soil Group. Front. Microbiol. 0, 208 (2012).

36. Wittig, I., Braun, H.-P. & Schägger, H. Blue native PAGE. (2006) doi:10.1038/nprot.2006.62.

37. Abby, S. S., Kerou, M. & Schleper, C. Ancestral Reconstructions Decipher Major Adaptations of Ammonia-Oxidizing Archaea upon Radiation into Moderate. MBio 11, (2020).

38. Rinke, C. et al. A standardized archaeal taxonomy for the Genome Taxonomy Database. Nat. Microbiol. 6, 946–959 (2021).

39. Abby, S. S. et al. Candidatus Nitrosocaldus cavascurensis, an ammonia oxidizing, extremely thermophilic archaeon with a highly mobile genome. Front. Microbiol. 9, 1–19 (2018).

40. Nicol, G. W. & Schleper, C. Ammonia-oxidising Crenarchaeota: important players in the nitrogen cycle? Trends Microbiol. 14, 207–212 (2006).

41. Luo, Z.-H. et al. Genomic Insights of “Candidatus Nitrosocaldaceae” Based on Nine New Metagenome-Assembled Genomes, Including “Candidatus Nitrosothermus” Gen Nov. and Two New Species of “Candidatus Nitrosocaldus”. Front. Microbiol. 11, (2021).

42. Hevler, J. F. et al. Selective cross-linking of coinciding protein assemblies by in-gel cross-linking mass spectrometry. EMBO J. (2021) doi:10.15252/embj.2020106174.

43. Stieglmeier, M. et al. Nitrososphaera viennensis gen. nov., sp. nov., an aerobic and mesophilic, ammonia-oxidizing archaeon from soil and a member of the archaeal phylum Thaumarchaeota. Int. J. Syst. Evol. Microbiol. 64, 2738–2752 (2014).

44. Albers, S. V. & Meyer, B. H. The archaeal cell envelope. Nat. Rev. Microbiol. 9, 414– 426 (2011).

45. Reyes, C. et al. Genome wide transcriptomic analysis of the soil ammonia oxidizing archaeon Nitrososphaera viennensis upon exposure to copper limitation. ISME J. 14, 2659–2674 (2020).

46. Qin, W. et al. Stress response of a marine ammonia-oxidizing archaeon informs physiological status of environmental populations. Nat. Publ. Gr. doi, (2017).

47. Carini, P., Dupont, C. L. & Santoro, A. E. Patterns of thaumarchaeal gene expression in culture and diverse marine environments. Environ. Microbiol. 20, 2112–2124 (2018).

48. Stewart, F. J., Ulloa, O. & Delong, E. F. Microbial metatranscriptomics in a permanent marine oxygen minimum zone. Environ. Microbiol. 14, 23–40 (2012).

49. Bayer, B. et al. Proteomic Response of Three Marine Ammonia-Oxidizing Archaea to Hydrogen Peroxide and Their Metabolic Interactions with a Heterotrophic Alphaproteobacterium Downloaded from. msystems.asm.org vol. 4 http://msystems.asm.org/ (2019).

50. Santoro, A. E. et al. Genomic and proteomic characterization of “Candidatus Nitrosopelagicus brevis”: an ammonia-oxidizing archaeon from the open ocean. Proc. Natl. Acad. Sci. U. S. A. 112, 1173–8 (2015).

51. Tolar, B. B. et al. Integrated structural biology and molecular ecology of N-cycling enzymes from ammonia-oxidizing archaea. Environ. Microbiol. Rep. 9, 484–491 (2017).

52. Diamond, S. et al. Soils and sediments host Thermoplasmata archaea encoding novel copper membrane monooxygenases (CuMMOs). ISME J. 1–15 (2022) doi:10.1038/s41396-021-01177-5.

53. Käll, L., Krogh, A. & Sonnhammer, E. L. L. A combined transmembrane topology and signal peptide prediction method. J. Mol. Biol. 338, 1027–1036 (2004).

54. Hakemian, A. S. & Rosenzweig, A. C. The Biochemistry of Methane Oxidation. (2007) doi:10.1146/annurev.biochem.76.061505.175355.

55. Jumper, J. et al. Highly accurate protein structure prediction with AlphaFold. Nature 596, (2021).

56. Varadi, M. et al. NAR Breakthrough Article AlphaFold Protein Structure Database : massively expanding the structural coverage of protein-sequence space with high-accuracy models. 50, 439–444 (2022).

57. Evans, R. et al. Protein complex prediction with AlphaFold-Multimer. bioRxiv (2021) doi:10.1007/978-1-61779-361-5_16.

58. Ernst, J. et al. STEM: a tool for the analysis of short time series gene expression data. BMC Bioinformatics 7, 191 (2006).

59. Klotz, M. G. & Stein, L. Y. Nitrifier genomics and evolution of the nitrogen cycle. (2007) doi:10.1111/j.1574-6968.2007.00970.x.

60. Lawton, T. J., Ham, J., Sun, T. & Rosenzweig, A. C. Structural conservation of the B subunit in the ammonia monooxygenase/particulate methane monooxygenase superfamily. Bone 23, 1–7 (2014).

61. Versantvoort, W. et al. Complexome analysis of the nitrite-dependent methanotroph Methylomirabilis lanthanidiphila. Biochim. Biophys. Acta -Bioenerg. 1860, 734–744 (2019).

62. Pitcher, A. et al. Crenarchaeol dominates the membrane lipids of Candidatus Nitrososphaera gargensis, a thermophilic Group I.1b Archaeon. ISME J. 4, 542–552 (2010).

63. Villanueva, L., Damsté, J.S.S. & Schouten, S. A re-evaluation of the archaeal membrane lipid biosynthetic pathway. Nat. Rev. Microbiol. 12, 438–448 (2014).

64. Sinninghe Damsté, J. S., Schouten, S., Hopmans, E. C., Van Duin, A. C. T. & Geenevasen, J. A. J. Crenarchaeol: The characteristic core glycerol dibiphytanyl glycerol tetraether membrane lipid of cosmopolitan pelagic crenarchaeota. J. Lipid Res. 43, 1641–1651 (2002).

65. De La Torre, J.R., Walker, C. B., Ingalls, A. E., Könneke, M. & Stahl, D. A. Cultivation of a thermophilic ammonia oxidizing archaeon synthesizing crenarchaeol. Environ. Microbiol. 10, 810–818 (2008).

66. Sinninghe Damsté, J. S. et al. Intact polar and core glycerol dibiphytanyl glycerol tetraether lipids of group I.1a and I.1b Thaumarchaeota in soil. Appl. Environ. Microbiol. 78, 6866–6874 (2012).

67. Elling, F. J., Könneke, M., Mußmann, M., Greve, A. & Hinrichs, K. U. Influence of temperature, pH, and salinity on membrane lipid composition and TEX86 of marine planktonic thaumarchaeal isolates. Geochim. Cosmochim. Acta 171, 238–255 (2015).

68. El Sheikh, A. F., Poret-Peterson, A. T. & Klotz, M. G. Characterization of two new genes, amoR and amoD, in the amo operon of the marine ammonia oxidizer Nitrosococcus oceani ATCC 19707. Appl. Environ. Microbiol. 74, 312–318 (2008).

69. Fisher, O. S. et al. Characterization of a long overlooked copper protein from methane-and ammonia-oxidizing bacteria. Nat. Commun. 9, 1–12 (2018).

70. Berube, P. M., Samudrala, R. & Stahl, D. A. Transcription of All amoC Copies Is Associated with Recovery of Nitrosomonas europaea from Ammonia Starvation. J. Bacteriol. 189, 3935–3944 (2007).

71. Berube, P. M. & Stahl, D. A. The divergent AmoC3 subunit of ammonia monooxygenase functions as part of a stress response system in Nitrosomonas europaea. J. Bacteriol. 194, 3448–56 (2012).

72. Lebedeva, E. V. et al. Enrichment and genome sequence of the group I.1a ammonia-oxidizing archaeon ‘Ca. Nitrosotenuis uzonensis’ representing a clade globally distributed in thermal habitats. PLoS One 8, 1–12 (2013).

73. Qin, W. et al. Nitrosopumilus maritimus gen. nov., sp. nov., Nitrosopumilus cobalaminigenes sp. nov., Nitrosopumilus oxyclinae sp. nov., and Nitrosopumilus ureiphilus sp. nov., four marine ammonia-oxidizing archaea of the phylum Thaumarchaeota. Int. J. Syst. Evol. Microbiol. (2017) doi:10.1099/ijsem.0.002416.

74. Ahlgren, N. A., Fuchsman, C. A., Rocap, G. & Fuhrman, J. A. Discovery of several novel, widespread, and ecologically distinct marine Thaumarchaeota viruses that encode amoC nitrification genes. ISME J. (2018) doi:10.1038/s41396-018-0289-4.

75. Hyman, M. R. & Wood, P. M. Methane oxidation by Nitrosomonas europaea. Biochem. J. 212, 31–37 (1983).

76. Hyman, M. R. & Wood, P. M. Ethylene oxidation by Nitrosomonas europaea. Arch. Microbiol. 137, 155–158 (1984).

77. Hyman, M. R., Murton, I. B. & Arp, D. J. Interaction of Ammonia Monooxygenase from Nitrosomonas europaea with Alkanes, Alkenes, and Alkynes. Appl. Environ. Microbiol. 54, 3187–3190 (1988).

78. Jones, R. D. & Morita, R. Y. Methane oxidation by Nitrosococcus oceanus and Nitrosomonas europaea. Appl. Environ. Microbiol. 45, 401–410 (1983).

79. Hyatt, D. et al. Prodigal: prokaryotic gene recognition and translation initiation site identification. BMC Bioinformatics 11, (2010).

80. Katoh, K., Misawa, K., Kuma, K. I. & Miyata, T. MAFFT: A novel method for rapid multiple sequence alignment based on fast Fourier transform. Nucleic Acids Res. 30, 3059–3066 (2002).

81. Katoh, K. & Standley, D. M. MAFFT multiple sequence alignment software version 7: Improvements in performance and usability. Mol. Biol. Evol. 30, 772–780 (2013).

82. Camacho, C. et al. BLAST+: Architecture and applications. BMC Bioinformatics 10, 1–9 (2009).

83. Criscuolo, A. & Gribaldo, S. BMGE (Block Mapping and Gathering with Entropy): A new software for selection of phylogenetic informative regions from multiple sequence alignments. BMC Evol. Biol. 10, (2010).

84. Minh, B. Q. et al. IQ-TREE 2: New Models and Efficient Methods for Phylogenetic Inference in the Genomic Era. Mol. Biol. Evol. 37, 1530–1534 (2020).

85. Hoang, D. T., Chernomor, O., Von Haeseler, A., Minh, B. Q. & Vinh, L. S. UFBoot2: Improving the ultrafast bootstrap approximation. Mol. Biol. Evol. 35, 518–522 (2018).

86. Graham, E. D., Heidelberg, J. F. & Tully, B. J. Potential for primary productivity in a globally-distributed bacterial phototroph. ISME J. 12, 1861–1866 (2018).

87. Rinke, C. et al. Insights into the phylogeny and coding potential of microbial dark matter. Nature 499, 431–437 (2013).

88. Tourna, M. et al. Nitrososphaera viennensis, an ammonia oxidizing archaeon from soil. Proc. Natl. Acad. Sci. USA 108, 8420–8425 (2011).

89. Reisinger, V. & Eichacker, L. A. Solubilization of membrane protein complexes for blue native PAGE. J. Proteomics 71, 277–283 (2008).

90. de Almeida, N. M. et al. Membrane-bound electron transport systems of an anammox bacterium: A complexome analysis. Biochim. Biophys. Acta - Bioenerg. 1857, 1694– 1704 (2016).

91. Berger, S., Cabrera-orefice, A., Jetten, M. S. M., Brandt, U. & Welte, C. U. Investigation of central energy metabolism-related protein complexes of ANME-2d methanotrophic archaea by complexome profiling. BBA - Bioenerg. 1862, 148308 (2021).

92. Schägger, H. & von Jagow, G. Tricine-sodium dodecyl sulfate-polyacrylamide gel electrophoresis for the separation of proteins in the range from 1 to 100 kDa. Anal. Biochem. 166, 368–379 (1987).

93. Schägger, H. Tricine-SDS-PAGE. Nat. Protoc. 1, (2006).

94. Rappsilber, J., Mann, M. & Ishihama, Y. Protocol for micro-purification, enrichment, pre-fractionation and storage of peptides for proteomics using StageTips. Nat. Protoc. 2, 1896–1906 (2007).

95. Tyanova, S., Temu, T. & Cox, J. The MaxQuant computational platform for mass spectrometry–based shotgun proteomics. Nat. Protoc. 11, (2016).

96. Pirklbauer, G. J. et al. MS Annika: A New Cross-Linking Search Engine. J. Proteome Res. 20, 2560–2569 (2021).

97. Iacobucci, C. et al. A cross-linking/mass spectrometry workflow based on MS-cleavable cross-linkers and the MeroX software for studying protein structures and protein–protein interactions. Nat. Protoc. 13, 2864–2889 (2018).

98. Bullock, J. M. A., Schwab, J., Thalassinos, K. & Topf, M. The importance of non-accessible crosslinks and solvent accessible surface distance in modeling proteins with restraints from crosslinking mass spectrometry. Mol. Cell. Proteomics 15, 2491–2500 (2016).

99. Kong, A. T., Leprevost, F. V., Avtonomov, D. M., Mellacheruvu, D. & Nesvizhskii, A. I. MSFragger: Ultrafast and comprehensive peptide identification in mass spectrometry-based proteomics. Nat. Methods 14, 513–520 (2017).

100. Kim, D., Paggi, J. M., Park, C., Bennett, C. & Salzberg, S. L. Graph-based genome alignment and genotyping with HISAT2 and HISAT-genotype. Nat. Biotechnol. 37, 907–915 (2019).

101. Liao, Y., Smyth, G. K. & Shi, W. FeatureCounts: An efficient general purpose program for assigning sequence reads to genomic features. Bioinformatics 30, 923–930 (2014).

102. Warnes, G. R. et al. gplots: Various R Programming Tools for Plotting Data. (2020).

103. R Core Team (2020). R: A language and environment for statistical computing. (2020).

104. Eren, A. M. et al. Community-led, integrated, reproducible multi-omics with anvi’o. Nat. Microbiol. 6, 3–6 (2021).

105. Capella-Gutiérrez, S., Silla-Martínez, J. M. & Gabaldón, T. trimAl: A tool for automated alignment trimming in large-scale phylogenetic analyses. Bioinformatics 25, 1972–1973 (2009).

106. Almagro Armenteros, J. J. et al. SignalP 5.0 improves signal peptide predictions using deep neural networks. Nat. Biotechnol. 37, 420–423 (2019).

107. Schrodinger LLC. The PyMOL Molecular Graphics System, Version 1.8. (2015).

108. Perez-Riverol, Y. et al. The PRIDE database and related tools and resources in 2019: Improving support for quantification data. Nucleic Acids Res. 47, D442–D450 (2019).

109. Larsson, A. AliView: A fast and lightweight alignment viewer and editor for large datasets. Bioinformatics 30, 3276–3278 (2014).

110. Berman, H. M. et al. The Protein Data Bank. Nucleic Acids Res. 28, 235–242 (2000).

