## Supplementary Material for "Unexpected Complexity of the Ammonia Monooxygenase in Archaea"

#### SDS-Tricine-PAGE verifies components of the AMO complex

To further explore the content of the AMO complex extracted from BN-PAGE, cut-outs from band 7 (Fig. 1) were subjected to SDS-Tricine-PAGE analysis under denaturing conditions [1,2]. SDS-PAGE bands containing AmoA, AmoB, AmoC, and AmoX were all identified using trypsin as the digestive enzyme (Supplementary Fig. 2). Apparent sizes of AmoB, AmoC, and AmoX proteins matched the expected molecular weight, while the band representing AmoA had a lower apparent molecular weight than expected, likely due to the high hydrophobicity of this subunit. Proteins with multiple transmembrane helices have been observed to run faster than expected in SDS gels [3]. Bands containing SlaA (S-layer protein), NVIE_028570 (exported protein of unknown function), and NVIE_024150 (ABC uptake transporter) were also identified. The band containing AmoB also contained a large amount of NVIE_021780 (exported protein of unknown function) and the one containing the highest amount for AmoX was dominated by the two hypothetical proteins identified from the syntenic analysis, NVIE_004540 and NVIE_004550, which are proposed here to be part of the archaeal AMO complex as AmoY and AmoZ respectively. The band representing AmoC could be separated into two protein groups: one representing amoC4/C6 that made up 96% of the total AmoC intensities, and one representing amoC1/C2 at 4%.

In an attempt to obtain unique AmoC peptides to distinguish AmoC4 and C6 which are 96 % identical in their amino acid sequence (BLASTp) [4,5], additional replicate cut-outs from BN-PAGE gels corresponding to band 7 were subjected to SDS-Tricine-PAGE. Bands corresponding to AmoC were digested using two different proteases (either GluC endoprotease or chymotrypsin). After chymotrypsin digestion it was possible to identify unique peptides from AmoC6, although the majority of AmoC peptides came from a shared region of both peptides. Peptide coverage maps for GluC, chymotrypsin, and trypsin digestion for the six homologs can be found in Supplementary Data. A unique peptide was not found for AmoC4 after correction of its predicted translational start site based on available transcriptomic and proteomic data (see details Methods). Therefore, it was not possible to definitively distinguish in these experiments between C4 and C6 as the dominant homolog in the AMO complex with proteomics alone.

**Architecture of the structural model of AmoABCXYZ in *N.viennensis* and *N. cavascurensis* and notes on the putative role of the novel subunits**

The Alphafold model of the *N.viennensis* AMO complex comprises single copies of the AmoA, AmoB, AmoC, AmoX, AmoY and AmoZ subunits (Fig. 3, Supplementary Fig. 8). Subunits AmoB and AmoZ contained predicted signal peptides which were not included in the modeled sequences. AmoA consists of six transmembrane (TM) helices and is encircled by the TM helices from AmoC, AmoB, AmoX and AmoZ, forming the core of the TM region of the complex. A loop and short helix formed between helices five and six protrude from the membrane and interact with the soluble domain from AmoB, as in the pMMO structure. AmoB consists of an N-terminal soluble β-barrel formed by seven antiparallel strands and a C-terminal TM helix which interacts with TM helices from AmoA and AmoZ . AmoC consists of four TM helices encircled by the helices of AmoA, AmoY and AmoX, and a pair of short C-terminal helices oriented perpendicular to the helices of AmoA on the cytoplasmic side of the molecule. AmoX consists of two TM helices and an N-terminal loop and short helix extending towards the cytoplasmic side of the complex, interacting putatively with the cytoplasmic section of AmoC. AmoY forms a single TM helix interacting with AmoC, with shorter helices extending towards both extracellular and cytoplasmic sides.

The proposed subunit AmoZ contains a soluble N-terminal domain formed by an extended disordered region and two alpha helices connected to a C-terminal transmembrane domain by an extended loop. The two alpha helices are stabilized by a disulfide bond between Cys 49 and Cys 59, conserved in the genus *Nitrososphaera* and in *Nitrosocaldaceae* (Fig. 3, Supplementary Fig. 8A,C,H). A hydrogen bond between Glu 48 and Arg 68 offers additional stabilization of the two helices (Supplementary Fig. 8H). Glu 48 is conserved in all AmoZ homologs, while the position of Arg 68 can also be occupied by a lysine or a glutamic acid. Interactions with the cupredoxin domain of AmoB can be observed in the form of hydrogen bonds between AmoZ-Arg 69 and AmoB-Glu 82 (Supplementary Fig. 8H). Both sites are universally conserved, corroborating the putative stabilizing role of the AmoZ. Interestingly, in *N. viennensis* Glu 61 and in *N. cavascurensis* Gln 60 are in close vicinity to the conserved AmoB histidines (His 125 and His 127 in *N. viennensis*) coordinating the $Cu_B$ metal site (Supplementary Fig. 8F), but it is unclear whether this affects the ionic environment of the metal site.

The $Cu_C$ copper site in NvAMO exhibits a few putative differences compared to the pMMO site. In addition to the canonical coordination residues Asp 41, His 45 and His 58 from AmoC, the conserved N-terminal histidine from AmoB (His30) is modelled as part of the coordination sphere, within 3Å of the previous residues (Supplementary Fig. 8E). In contrast, the amino-terminal histidine is part of the $Cu_B$ centre in pMMO. In NcavAMO, the flexibility of the N-terminal region of AmoB could result in the same orientation. Interestingly, the N-terminal tyrosine of AmoZ (Tyr 31) in NvAMO is modelled in close vicinity (within 4.5-7.2 Å) to the $Cu_C$ coordination sphere (Fig S8E). Since this residue is universally conserved, implying an essential role, and the N-terminus of AmoZ is a flexible region, it is interesting to speculate whether it can influence the charge distribution in the vicinity of this metal site in archaea.

Regarding the novel subunits AmoY and AmoX, in addition to the stabilizing role of their TM helices for the holoenzyme, they both encode cytoplasmic helices which could be important for protein interactions as part of signaling cascades in the cytoplasmic side.

The predicted models of NvAMO and NcavAMO are similar in their general architecture, with a few notable differences (Supplementary Fig S1D). The relative positioning of AmoZ is discussed in the main text, and due to its overall weaker interactions with the complex it is evident that high-confidence modeling of this subunit is not possible with current methods, shown also by its overall lower confidence score in the models (Supplementary Fig. 8B,D). NcavAmoB encodes an extended section of the N-terminal extracellular domain between β strands 1 and 2 (residues 55-81), which forms two short helices and two loops and is anchored to the core β-barrel domain with a disulfide bond between Cys 76 and Cys 140 and multiple hydrogen bonds (Supplementary Fig. 8C,G), as already observed by Lawton et al. (2014) [6] in the crystal structure of AmoB from *Ca.* Nitrosocaldus yellowstonensis. The putative role of this extension could be an enhanced stability of the enzyme in the high temperatures experienced by this lineage.

Superimposition of the NvAMO model to the crystal structure of the particulate methane monooxygenase (PDB:1YEW, pMMO) from Methylococcus capsulatus (Bath) (B), reveals conservation of the overall fold of the main subunits AmoA, AmoB and AmoC, with an RMSD of 5.544 Å and 1573 atoms aligned (Supplementary Fig. 7A,B,C). Known differences include the absence of the second cupredoxin domain and the second TM helix of PmoB in AmoB, absence of the seventh TM helix of PmoA in AmoA, and the different localization of AmoC in the multimer compared to PmoC, which in part accommodates the two TM helices of AmoX.

### References


1. Schägger, H. & von Jagow, G. Tricine-sodium dodecyl sulfate-polyacrylamide gel electrophoresis for the separation of proteins in the range from 1 to 100 kDa. *Anal. Biochem.* **166**, 368–379 (1987).
2. Schägger, H. Tricine-SDS-PAGE. *Nat. Protoc.* **1**, (2006).
3. Rath, A. & Deber, C. M. Correction factors for membrane protein molecular weight readouts on sodium dodecyl sulfate-polyacrylamide gel electrophoresis. (2012) doi:10.1016/j.ab.2012.11.007.
4. Altschul, S. F. *et al.* Protein database searches using compositionally adjusted substitution matrices. *FEBS J.* **272**, 5101–5109 (2005).
5. Altschul, S. F. *et al.* Gapped BLAST and PSI-BLAST: a new generation of protein database search programs. *Nucleic Acids Res.* **25**, 3389–3402 (1997).
6. Lawton, T. J., Ham, J., Sun, T. & Rosenzweig, A. C. Structural conservation of the B subunit in the ammonia monooxygenase/particulate methane monooxygenase superfamily. *Bone* **23**, 1–7 (2014).

### Supplementary Figures

**A**

**B**

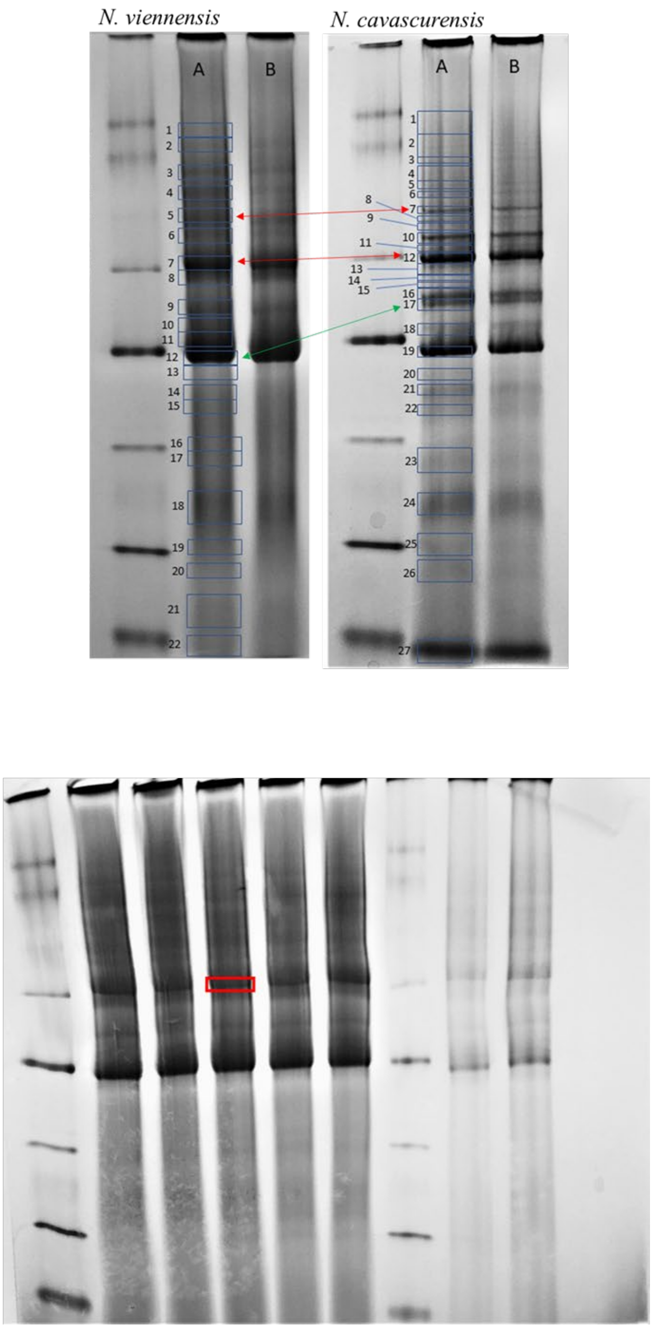


**Supplementary Figure 1. BN-PAGE gels of *N. viennensis* and *N.* cavascurensis. A.)** Comparison of protein membrane fractions between two species on BN-PAGE gels. Ladder and gels are the same. Blue boxes represent cut bands for proteomic analysis. Red arrows indicate bands of high AMO content that correspond between the two species. Green arrow points to a third AMO peak in both species that is found at different heights. A and B represent identical gel and samples conditions in each gel. **B.)** BN-PAGE gel of *N. viennensis* used for in-gel cross linking. The red box indicates the cut-out used for in-gel cross-linking with DSSO and subsequent mass spectrometry analysis. This cut-out corresponds to band 7 within Figure 1.

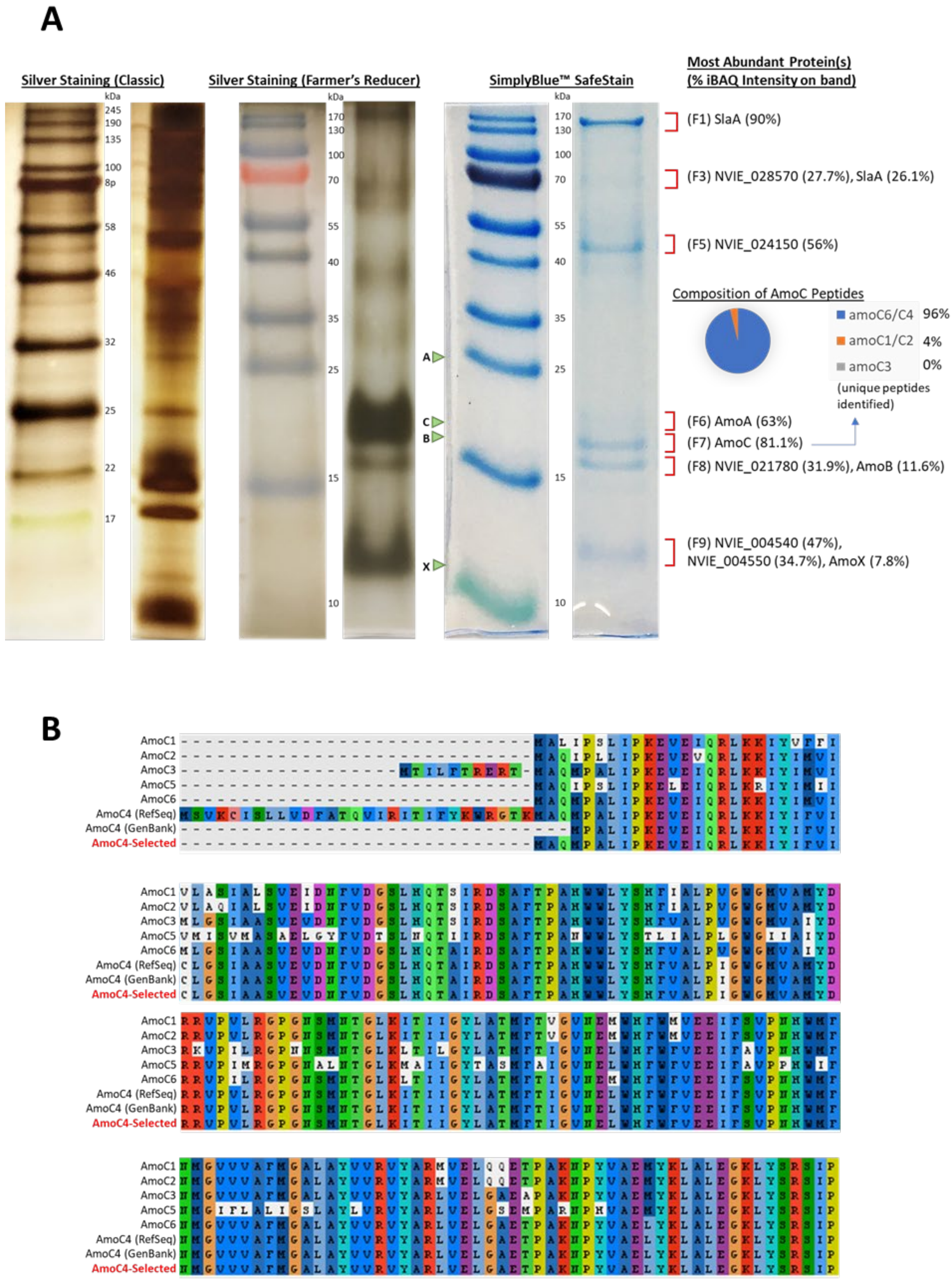


**Supplementary Figure 2. SDS-Tricine-PAGE gels of AMO cut-outs and AmoC homolog alignments from *N. viennensis*. A.)** SDS-Tricine-PAGE of AMO bands from BN-PAGE gels. Comparison of three different staining methods for SDS-Tricine-PAGE gels with size markers run on left side. Bands cut for analysis from a gel stained with SimplyBlue SafeStain and digested using trypsin are indicated by red brackets. Percentages represent the percentage of iBAQ normalized protein intensities for each individual band. Band identifiers are indicated in parentheses. Green arrows marked A, B, C, and X represent expected heights of bands for AMO subunits AmoA, AmoB, AmoC, and AmoX respectively. A pie chart from the band with the highest amount of AmoC shows the percentage of AmoC bands coming from different distinguished AmoC homologs. **B.)** Alignment of AmoC homologs from *N. viennensis*. Annotation differences can be seen for the AmoC4 homolog. The annotation marked in red font was chosen for analysis of AmoC peptides. Alignment created using AliView 1.27 [109].

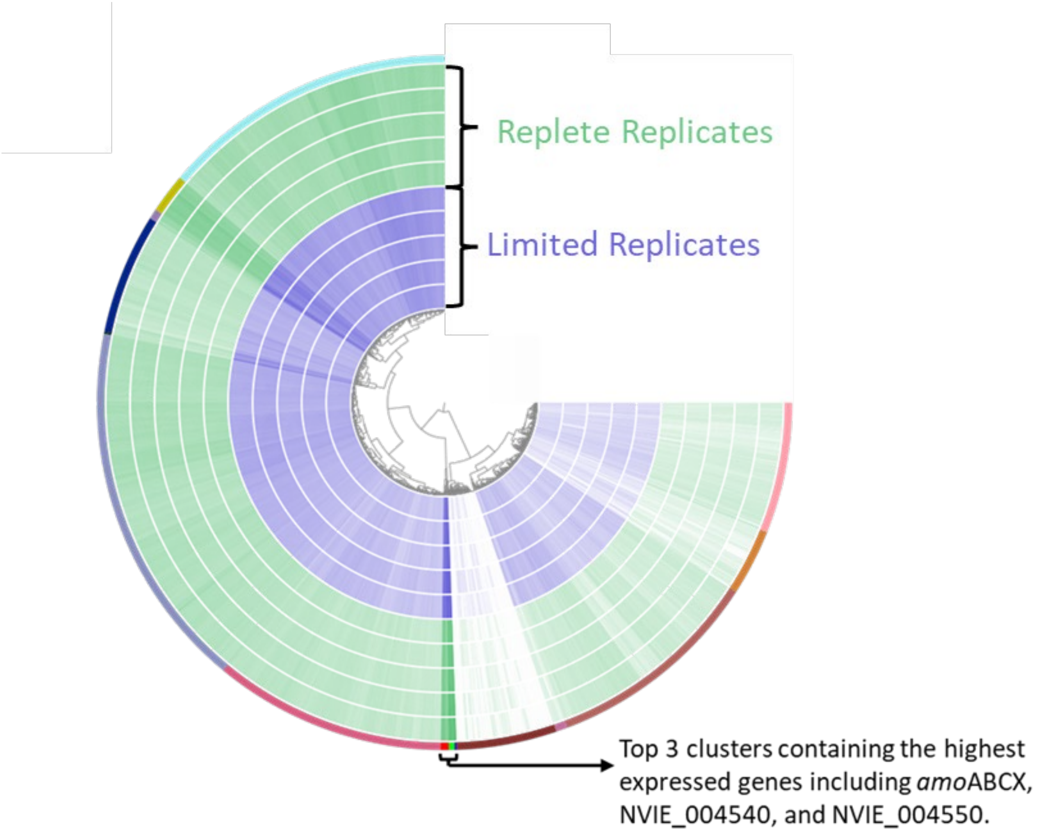


**Supplementary Figure 3. Transcriptomic clustering analysis of AMO subunits in *N. viennensis*. A.)** Clustering analysis of gene expression in *N. viennensis* under copper replete and limited conditions from Reyes et al. (2020)[45]. Genes were clustered into 15 separate clusters represented by different colors on the outer edge of the circular plot.

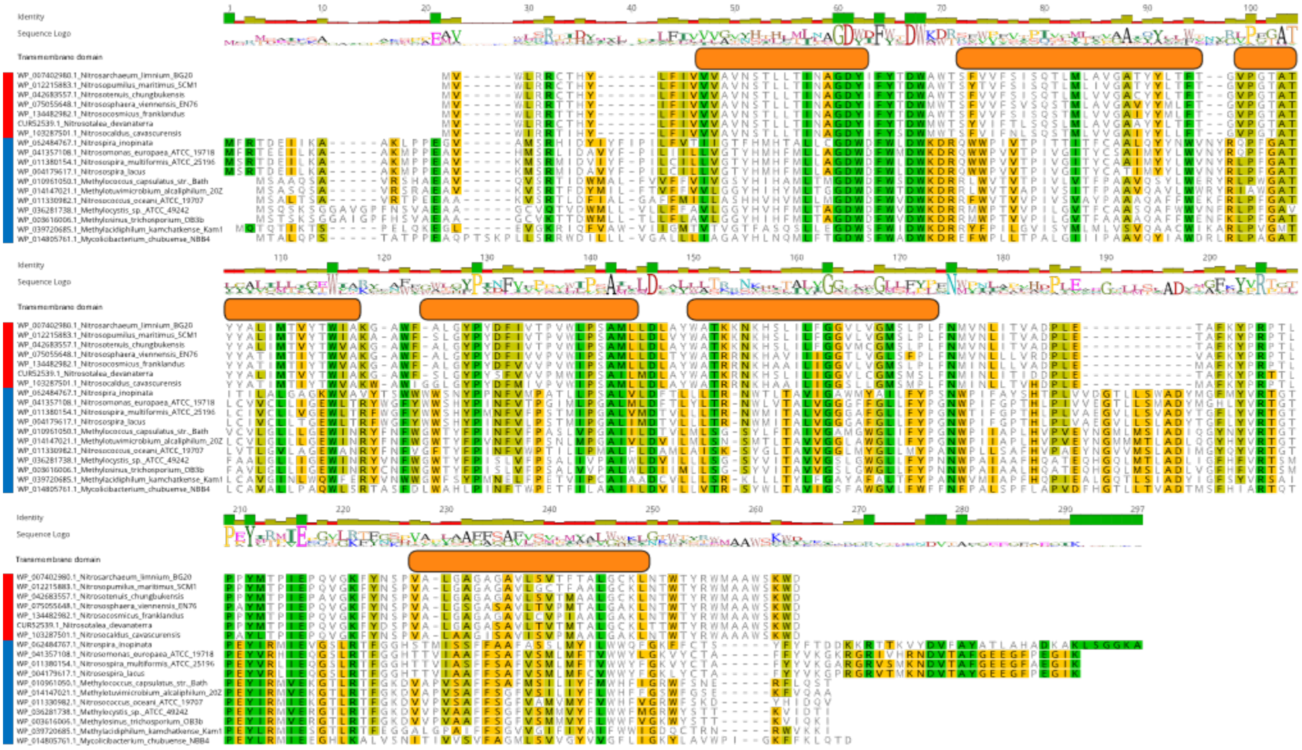


**Supplementary Figure 4. Protein alignments of PmoA/AmoA subunits from selected species.** Percent identity of amino acids across species are indicated. Sequence logo visually represents the proportion of amino acids at each position. Orange bars represent transmembrane helices as found in the crystal structure of PMO from *Methylosinus trichosporium* OB3b [23]. Red bars represent archaeal species and blue bars represent bacterial species.

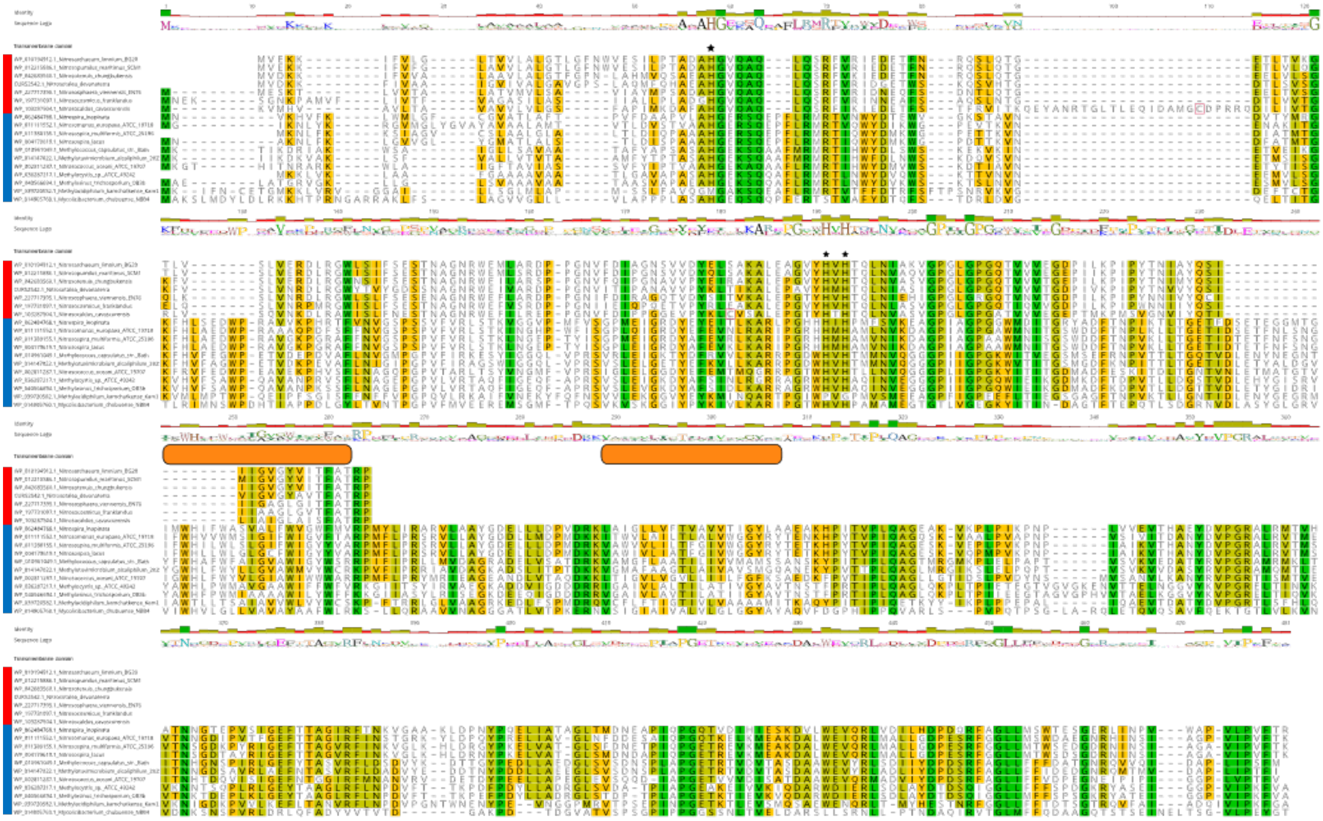

**Supplementary Figure 5. Protein alignments of PmoB/AmoB subunits from selected species.** Percent identity of amino acids across species is indicated. Sequence logo visually represents the proportion of amino acids at each position. Orange bars represent transmembrane helices as found in the crystal structure of PMO from *Methylosinus trichosporium* OB3b [23]. Red bars represent archaeal species and blue bars represent bacterial species. Conserved metal binding residues (with the exception of *Methylacidphilum kamchatkense* Kam1 from Verrucomicrobia) are marked with a black star.

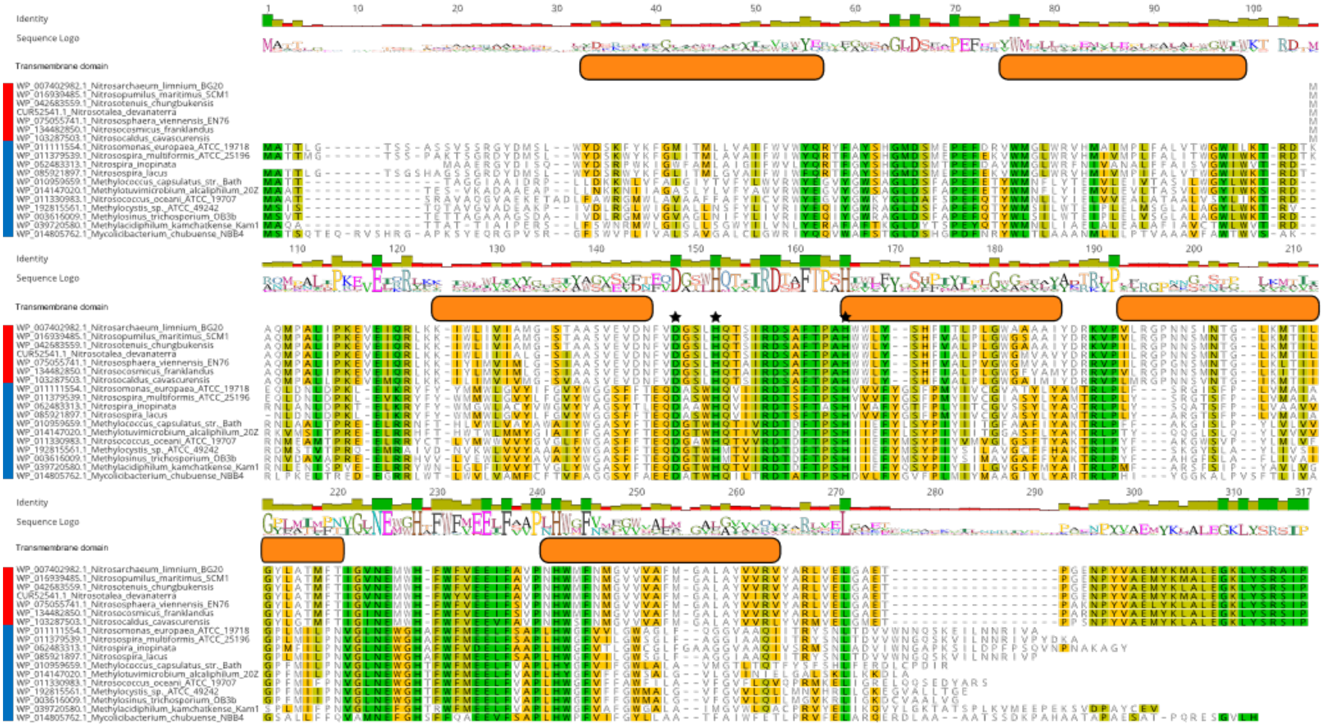


**Supplementary Figure 6. Protein alignments of PmoC/AmoC subunits from selected species.** Percent identities of amino acids across species is indicated. Sequence logo visually represents the proportion of amino acids at each position. Orange bars represent transmembrane helices as found in the crystal structure of PMO from *Methylosinus trichosporium* OB3b [23]. Red bars represent archaeal species and blue bars represent bacterial species. Conserved metal binding residues are marked with a black star.

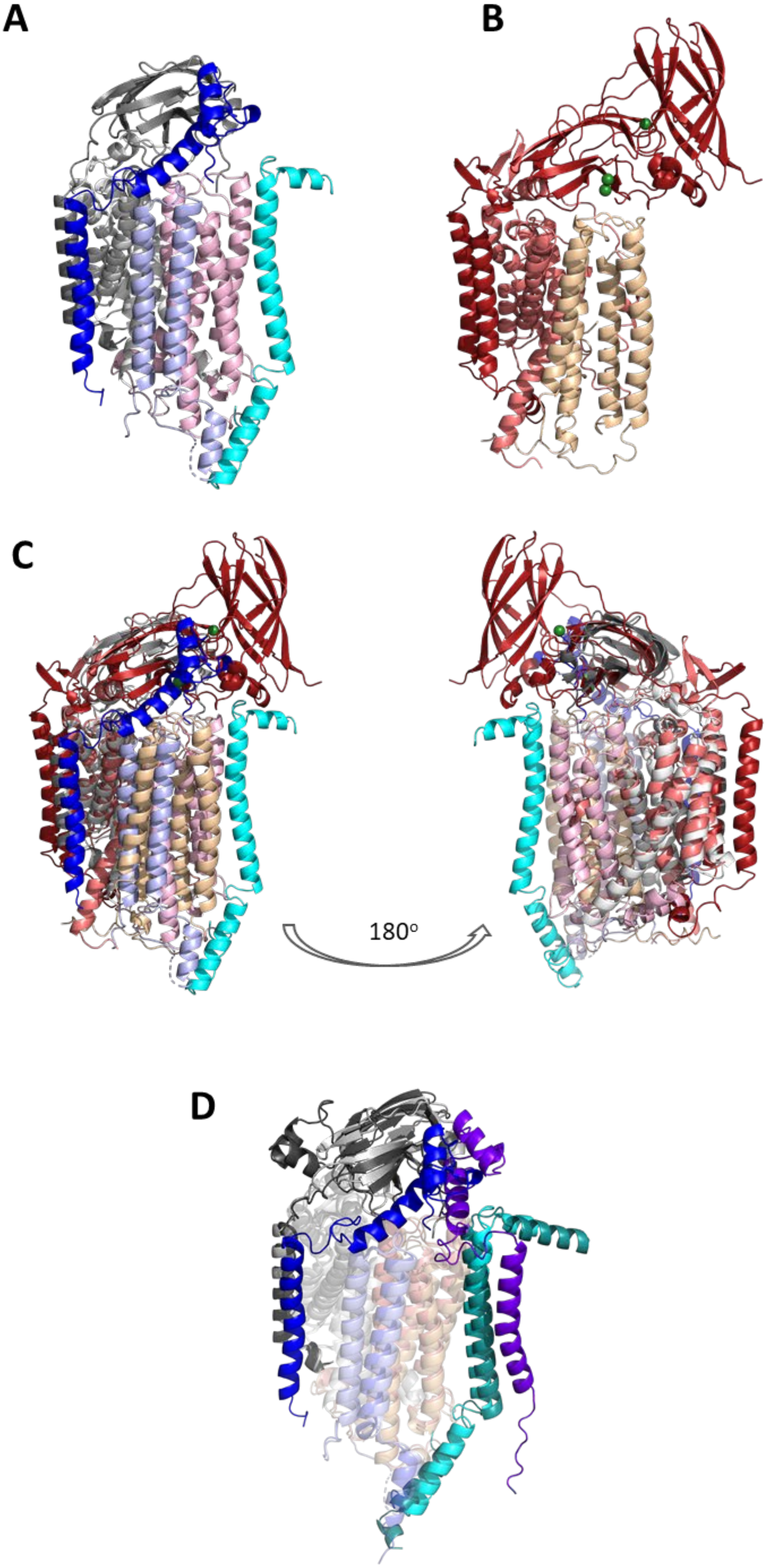
A
B
C
180°
D

**Supplementary Figure 7. Structural comparison between archaeal AMO and bacterial PMO. A.)** Alphafold model of NvAmoABCXYZ. **B.)** Crystal structure of the particulate methane monooxygenase (PDB:1YEW, pMMO) from Methylococcus capsulatus (Bath) [22] downloaded from the Protein Data Bank (PDB) [110]. **C.)** Superimposition of the Alphafold model of NvAmoABCXYZ **(A)** on the crystal structure of the particulate methane monooxygenase (PDB:1YEW, pMMO) from Methylococcus capsulatus (Bath) **(B)**, with an RMSD 5.544 Å. Subunits are colored as follows: NvAmoA, light grey; NvAmoB, dark grey; NvAmoC, light pink; NvAmoX, light blue; NvAmoY, cyan; NvAmoZ, blue; PmoB, dark red; PmoA, salmon; PmoC, sand. **D.)** Superimposition of the Alphafold model of NcavAmoABCXYZ on the NvAmoAMBXYZ, with an RMSD of 1.67Å, side view. Only subunits that exhibit substantial differences are in full colour: NvAmoB, light grey; NcavAmoB, dark grey; NvAmoY, cyan; NcavAmoY, teal; NvAmoZ, light blue; NcavAmoZ, purple. Subunits AmoA, AmoC and AmoX are in transparent tones of grey, salmon and blue respectively (dark for N.cavascurensis and light for N. viennensis). Images generated using PyMol[107].

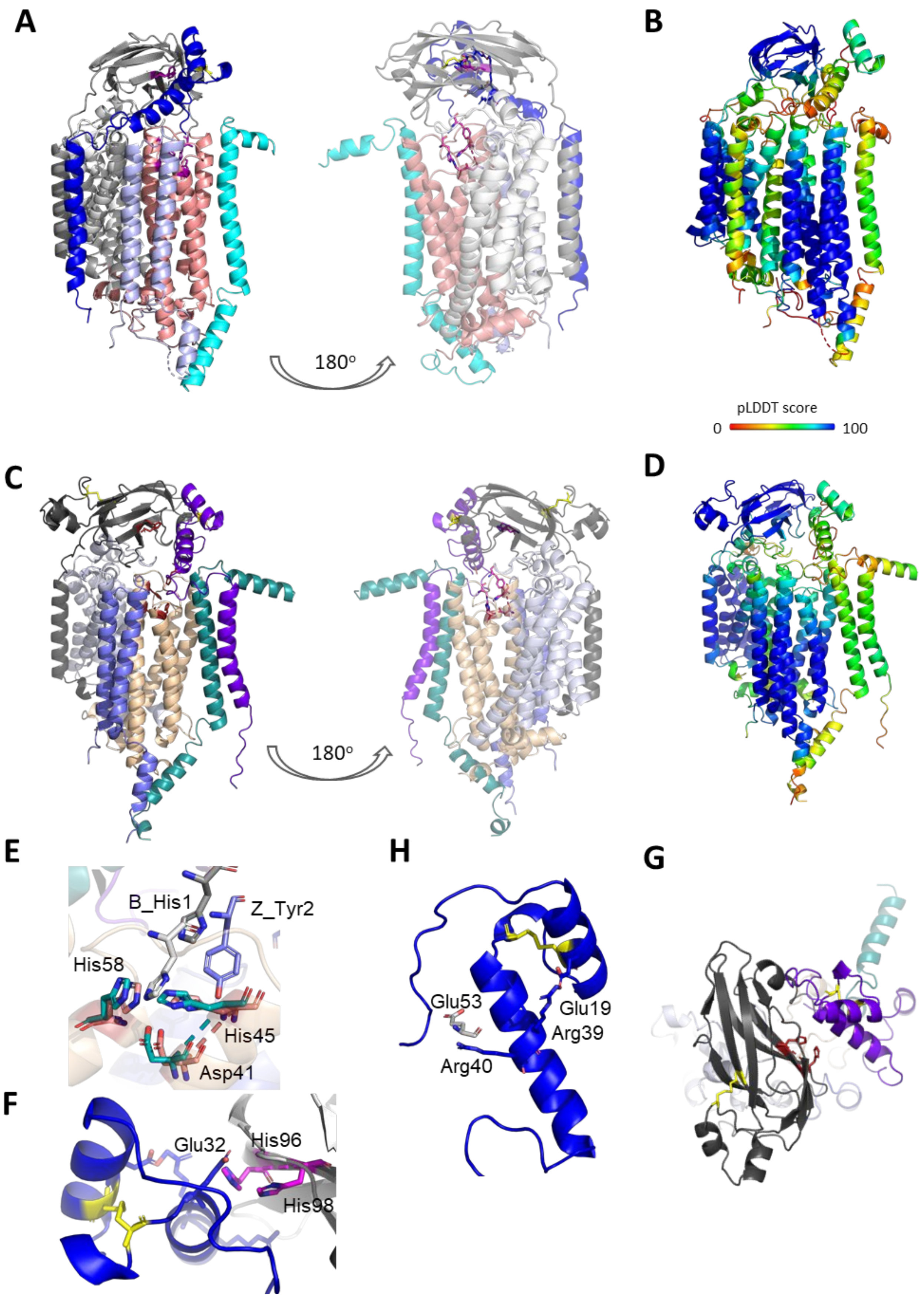
A
180°
B
pLDDT score
0
100
C
180°
D
E
B_His1
Z_Tyr2
His58
His45
Asp41
F
Glu32
His96
His98
H
Glu53
Glu19
Arg39
Arg40
G

**Supplementary Figure 8. Features of the AlphaFold models of NvAMO and NcavAMO heterohexamers. A, C.)** Different views of the NvAmoABCXYZ hexamer **(A)** and the NcavAmoABCXYZ hexamer **(C)** in cartoon representations, revealing the position of the $Cu_B$ and $Cu_C$ copper centers in magenta sticks, and the disulfide bonds in yellow. Subunits are colored as follows: NvAmoA, light grey; NvAmoB, grey; NvAmoC, salmon; NvAmoX, light blue; NvAmoY, cyan; NvAmoZ, blue; NcAmoA, light grey; NcAmoB, dark grey; NcAmoC, sand; NcAmoX, sky blue; NcAmoY, teal; NcAmoZ, purple. **B, D.)** The NvAmoABCXYZ and NcavAmoABCXYZ models colored according to the per residue pLDDT confidence scores from red (0) to blue (100). 57% of the residues in NvAmoABCXYZ and 52% in NcavAmoABCXYZ were modeled with a relatively high accuracy of >70%. **E.)** Closeup view of the residues involved in the coordination of the $Cu_C$ copper center. Canonical residues His58, His45, Asp41 from AmoC (*N.viennensis* numbering) are colored in salmon for the NvAMO model and teal for the NcavAMO model respectively. The conserved His30 from the N-terminus of AmoB which might be participating in the coordination sphere is depicted in white and grey for NvAMO and NcavAMO respectively. The conserved Tyr31 from NvAmoZ also located in the vicinity (4.5-7.2 Å) of the metal coordination sphere is depicted in blue. **F.)** Closeup view of the residues involved in the coordination of the NvAMO $Cu_B$ copper center. Canonical residues His125 and His127 (AmoB) are depicted in magenta, while Glu61 from AmoZ located in the close vicinity of the center is in blue. **H.)** Closeup view of the residues involved in putatively stabilizing intra- and inter-subunit interactions between AmoZ (blue) and AmoB (grey). A hydrogen bond between conserved residues Glu48 and Arg68 could stabilize the extracellular helices of AmoZ in addition to the predicted disulfide bond (in yellow), while conserved Arg69 could form a hydrogen bond with Glu82 from AmoB. **G.)** Top view of the extracellular domain in the NcavAMO model, illustrating the relative orientation of AmoB (grey) and AmoZ (purple). Note the disulfide bonds (in yellow) tethering the extended AmoB loop to the β-barrel, and between the AmoZ extracellular helices. The $Cu_B$ center is depicted in dark red sticks.

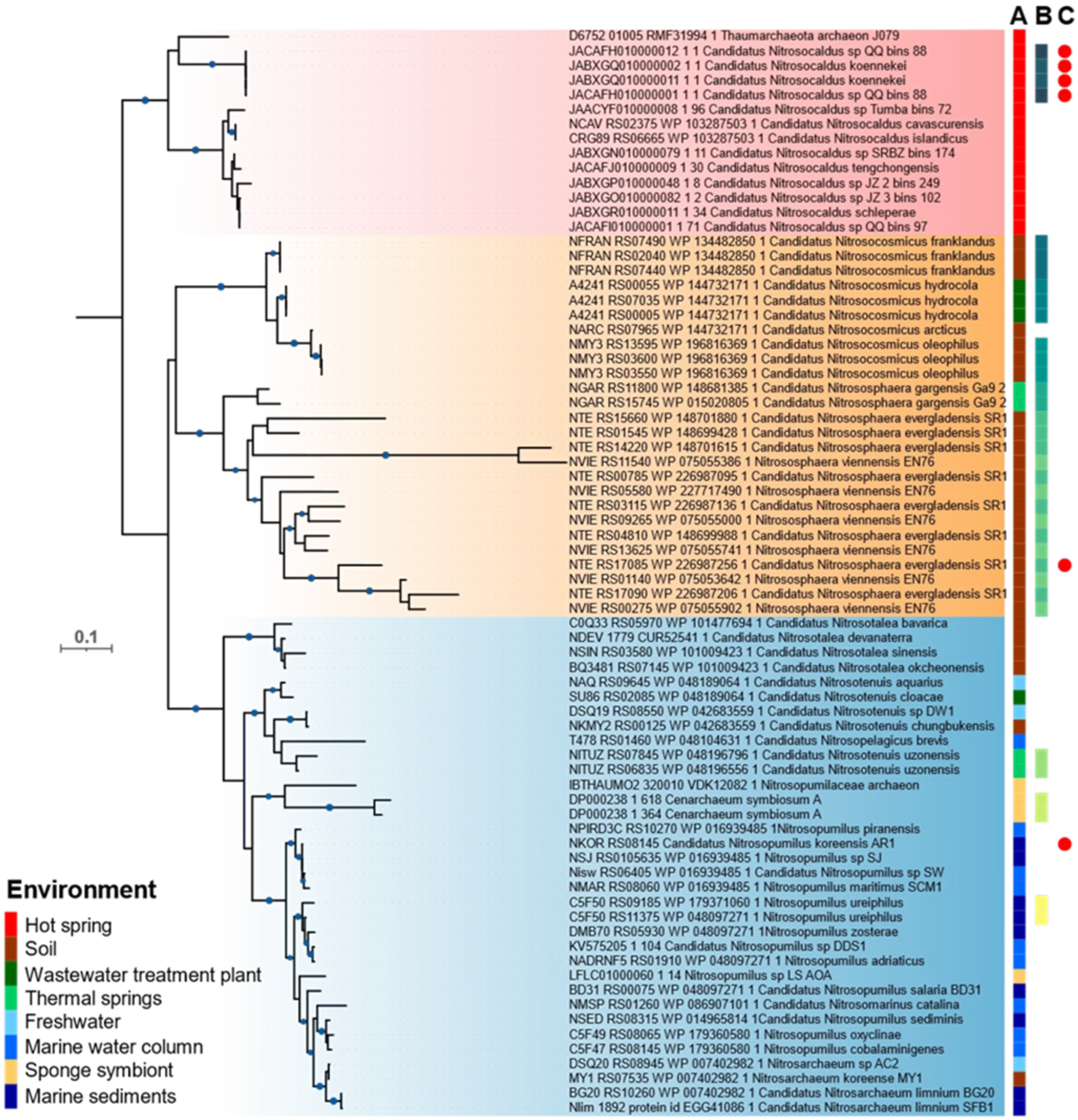


**Supplementary Figure 9. Phylogenetic tree of *amo*C gene nucleotide sequences in ammonia oxidizing archaea.** Labels are colored according to GTDB[38] families: *Nitrosocaldaceae*-red, *Nitrosopumilaceae*-blue, *Nitrososphaeraceae*-orange. Blue circles represent bootstrap values ≥85%. Labels include the gene locus tag followed by the NCBI protein accession and species name. Duplications are colored according to species. Added information next to the phylogenetic tree includes: (A) environment; (B) *amo*C duplications represented by different colors for each specie; (C) the *amo*C shown is a fragment (full circle).

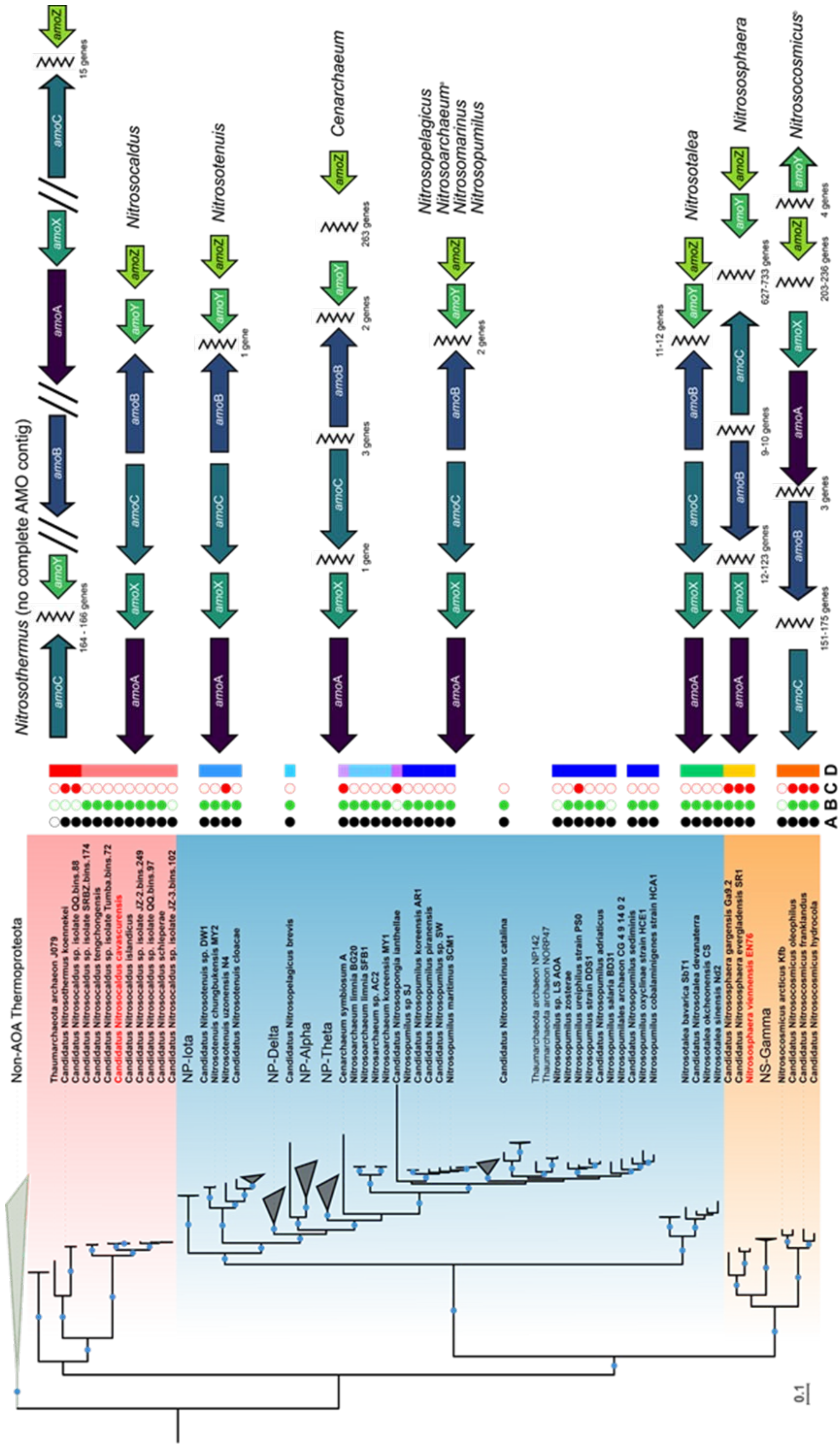
Nitrosothermus (no complete AMO contig)
Nitrosocaldus
Nitrosotenuis
Cenarchaeum
Nitrosopelagicus
Nitrosoarchaeum
Nitrosomarinus
Nitrosopumilus
Nitrosotalea
Nitrosospaera
Nitrosocosmicus
amoA
amoB
amoC
amoX
amoY
amoZ
164 - 166 genes
15 genes
1 gene
3 genes
2 genes
263 genes
11-12 genes
9-10 genes
627-733 genes
12-123 genes
151-175 genes
203-236 genes
4 genes
Non-AOA Thermoproteota
Thaumarchaeota archaeon J079
Candidatus Nitrosothermus koennekei
Candidatus Nitrosocaldus sp. isolate QQ.bins.88
Candidatus Nitrosocaldus sp. isolate SRBZ.bins.174
Candidatus Nitrosocaldus tengchongensis
Candidatus Nitrosocaldus sp. isolate Tumba.bins.72
Candidatus Nitrosocaldus cavascurensis
Candidatus Nitrosocaldus islandicus
Candidatus Nitrosocaldus sp. isolate JZ-2.bins.249
Candidatus Nitrosocaldus sp. isolate QQ.bins.97
Candidatus Nitrosocaldus schieperae
Candidatus Nitrosocaldus sp. isolate JZ-3.bins.102
NP-Iota
Candidatus Nitrosotenuis sp. DW1
Candidatus Nitrosotenuis chungbukensis MY2
Nitrosotenuis uzonensis N4
Candidatus Nitrosotenuis cloacae
NP-Delta
Candidatus Nitrosopelagicus brevis
NP-Alpha
NP-Theta
Cenarchaeum symbiosum A
Nitrosoarchaeum limnia BG20
Nitrosoarchaeum limnia SFB1
Nitrosoarchaeum sp. AC2
Nitrosoarchaeum koreensis MY1
Candidatus Nitrosospongia ianthellae
Nitrosopumilus sp SJ
Candidatus Nitrosopumilus koreensis AR1
Candidatus Nitrosopumilus piranensis
Candidatus Nitrosopumilus sp. SW
Nitrosopumilus maritimus SCM1
Candidatus Nitrosomarinus catalina
Thaumarchaeota archaeon NP142
Thaumarchaeota archaeon NORP47
Nitrosopumilus sp. LS AOA
Nitrosopumilus zosterae
Nitrosopumilus ureiphilus strain PS0
Nitrosopumilus strain DDS1
Candidatus Nitrosopumilus adriaticus
Candidatus Nitrosopumilus salaria BD31
Nitrosopumilales archaeon CG 4 9 14 0 2
Candidatus Nitrosopumilus sediminis
Nitrosopumilus oxyclinae strain HCE1
Nitrosopumilus cobalaminigenes strain HCA1
Nitrosotalea bavarica SbT1
Candidatus Nitrosotalea devanaterra
Nitrosotalea okcheonensis CS
Nitrosotalea sinensis Nd2
Candidatus Nitrosphaera gargensis Ga9.2
Candidatus Nitrosphaera evergladensis SR1
Nitrososphaera viennensis EN76
NS-Gamma
Nitrosocosmicus arcticus Kfb
Candidatus Nitrosocosmicus oleophilus
Candidatus Nitrosocosmicus franklandus
Candidatus Nitrosocosmicus hydrocola
A B C D
0.1

**Supplementary Figure 10. Genomic comparison of AMO subunit synteny in AOA (extended version of Figure 2).** The left side of the figure represents a phylogenetic tree of AOA based on 32 conserved ribosomal proteins. Blue circles represent bootstrap values of 100%. Taxonomic labels are colored according to GTDB family identity: *Nitrosocaldaceae*-red, *Nitrosopumilaceae*-blue, *Nitrososphaeraceae*-orange. Labels in bold were included in syntenic analysis. Labels in red represent species with proteomic evidence from BN-PAGE gels. Homologs of NVIE_004540 are represented by AmoY and homologs of AmoZ are represented by AmoZ. Separate contigs are represented by a double forward slash. Gaps between genes on the same contig are marked by a zig-zag line. Added information next to the phylogenetic tree includes: (A) all six AMO subunits found (full circle); (B) AMO subunits found on the same contig (full circle); (C) multiple *amo*C copies found within the genome (full circle); (D) different genera indicated by different colors: *Nitrosothermus* (bright red), *Nitrosocaldus* (pink), *Nitrosotenuis* (sea blue), *Nitrosopelagicus* (aqua), *Cenarchaeaum* (light purple), *Nitrosoarchaeum* (light blue), *Nitrosospongia* (dark purple), *Nitrosopumilus* (dark blue), *Nitrosotalea* (green), *Nitrosocosmicus* (yellow), *Nitrososphaera* (orange). With the exception of *Nitrosothermus* sp., species with AMO genes on different contigs could still plausibly fit into the shown arrangements. The sponge species *Ca.* Nitrospongia ianthellae and *Ca*. Nitrosopumilus sp. LS AOA, as well as *Nitrosoarchaeum limnia* SFB1 differ from the other arrangements of marine AOA by having 1 gene inserted between AmoY and AmoZ rather than 2.
